# Engineering and Evolution of the Complete Reductive Glycine Pathway in *Saccharomyces cerevisiae* for Formate and CO_2_ Assimilation

**DOI:** 10.1101/2023.07.10.548313

**Authors:** Viswanada R Bysani, Ayesha S M Alam, Arren Bar-Even, Fabian Machens

## Abstract

Using captured CO_2_ and C1-feedstocks like formate and methanol derived from electrochemical activation of CO_2_ are key solutions for transforming industrial processes towards a circular carbon economy. Engineering formate and CO_2_-based growth in the biotechnologically relevant yeast *Saccharomyces cerevisiae* could boost the emergence of a formate-mediated circular bio-economy. This study adopts a growth-coupled selection scheme for modular implementation of the Reductive Glycine Pathway (RGP) and subsequent Adaptive Laboratory Evolution (ALE) to enable formate and CO_2_ assimilation for biomass formation in yeast. We first constructed a serine biosensor strain and then implemented the serine synthesis module of the RGP into yeast, establishing glycine and serine synthesis from formate and CO_2_. ALE improved the RGP-dependent growth by 8-fold. 13C-labeling experiments reveal glycine, serine, and pyruvate synthesis via the RGP, demonstrating the complete pathway activity. Further, we reestablished formate and CO_2_-dependent growth in non-evolved biosensor strains via reverse-engineering a mutation in *GDH1* identified from ALE. This mutation led to significantly more 13C-formate assimilation than in WT without any selection or overexpression of the RGP. Overall, we demonstrated the activity of the complete RGP, showing evidence for carbon transfer from formate to pyruvate coupled with CO_2_ assimilation.

## Introduction

The enormous dependency on finite fossil resources for energy and commodity products is unsustainable and led to increased CO_2_ and other greenhouse gas emissions since the Industrial Revolution, causing global warming^1^. Using agriculture-based renewable sugar substrates and lignocellulosic feedstocks for bioprocesses competes with food production and requires intensive land use, making these approaches less sustainable and morally questionable^2^. The progressing climate catastrophe and a growing world population force humankind to shift rapidly towards a sustainable bioeconomy using suitable renewable substrates, ideally including circularization of CO_2_ emission. A recent update on *our world in data* shows that >35 bt of CO_2_ were released into the atmosphere alone in 2021^3^ without signs of stabilization. Using CO_2_ captured from industrial points or the atmosphere as the substrate for manufacturing while shifting towards sustainable energy usage offers a solution to mitigate climate change.

Various CO_2_ capture technologies have emerged in recent years, aiming to reuse CO_2_ directly or indirectly. These methods include the efficient (electro)chemical conversion of CO_2_ into mediator one-carbon (C1) compounds formate^4–6^ and methanol^7,8^ using renewable energy^9,10^. These reduced C1 compounds are promising alternative feedstocks to transform sugar-based bioprocessing because they can be produced from captured CO_2_ at high energy conversion efficiency, and are close to commercialization^11^. Here, we focus on formate, which can be sourced through the direct reduction of CO_2_ in a single electro-chemical system.

However, not all microbes are able to grow on formate or other one-carbon molecules as a feedstock, and those microbes that are, are generally not well-established for industrial processes^12,13^. Thus, the latter are not attractive hosts for a formate-based bio-economy. Engineering industrially-relevant organisms, such as *Saccharomyces cerevisiae*, *Escherichia coli*, and others, is thus a promising approach. We previously determined the synthetic RGP as the most promising, oxygen-tolerant pathway to establish synthetic formatotrophy^14–17^.

Briefly, the RGP consists of four modules. A glycine biosynthesis module, in which formate is attached to the C1-carrier tetrahydrofolate (THF), reduced to CH_2_-THF, and then, by the reversed activity of the glycine cleavage system (rGCS), condensed with ammonium and CO_2_ to yield glycine. Next, the glycine-to-serine conversion module condenses glycine and hydroxymethyl from formate-derived CH_2_-THF to produce serine. Finally, the pyruvate synthesis module converts serine into pyruvate by deamination. Pyruvate then feeds into the central metabolic network and eventually can support full cell growth. The accessory energy module, composed of a formate dehydrogenase, provides the required energy and reducing power generated by oxidation of formate to CO_2_ and NADH. Our group has successfully engineered formatotrophic *E. coli* and *Cupriavidus necator* strains for growth via the RGP^16,17^. A few recent studies show that this pathway is also operating naturally in several specialized prokaryotes^18–21^. Nevertheless, these microbes are not well-established for industrial applications, as they are, e.g., slow growing, strictly anaerobic, and genetically not accessible yet. *S. cerevisiae* is an organism generally regarded as safe (GRAS) and well-studied for several industrial applications^22,23^. Thus, establishing formatotrophic growth in yeast is a key goal for establishing a C1-based bioeconomy. Despite its vast potential for industrial applications, yeast is especially interesting for engineered formatotrophy since it can tolerate very high levels of formate and grow in low pH environments^24^. The latter opens the perspective easier feeding strategies using protonated formic acid rather than formic acid salt, e.g., sodium formate, the latter will require acid addition to control pH. Further, baker’s yeast has the advantage that all enzymes required to operate the RGP are endogenous; thus, the expression of foreign genes is not strictly required. Hence, we aimed to engineer the RGP in *S. cerevisiae*.

The yeast enzymes necessary for the glycine synthesis module are the trifunctional Mis1p, which catalyzes formate ligation with THF and the subsequent reduction of CHO-THF to CH_2_-THF, and Gcv1p, Gcv2p, Gcv3p, and Lpd1p - together forming the GSC (Glycine Synthase Complex). Gcv3p (GcvH) acts as the core shuttle protein in the GSC and binds the lipoic acid cofactor. Lpd1p reduces the lipoyl moiety bound to Gcv3p, using NADH as a cofactor. Gcv1p (GcvT) then condenses one molecule of CH_2_-THF with one molecule of NH_3_ on the reduced lipoyl moiety, thereby releasing a THF.

The Gcv2p (GcvP), with a pyridoxal-5-phosphate as a cofactor, then adds one molecule of CO_2_ to the aminomethyl group on the lipolyl moiety releasing glycine from Gcv3p. Finally, the oxidized lipoic acid bound to Gcv3p is reduced by Lpd1p to allow the next glycine synthesis cycle (**Figure 1**). This core module requires one molecule of ATP, NADH, and NADPH for each molecule of formate that is assimilated into glycine.

**Figure 1:**
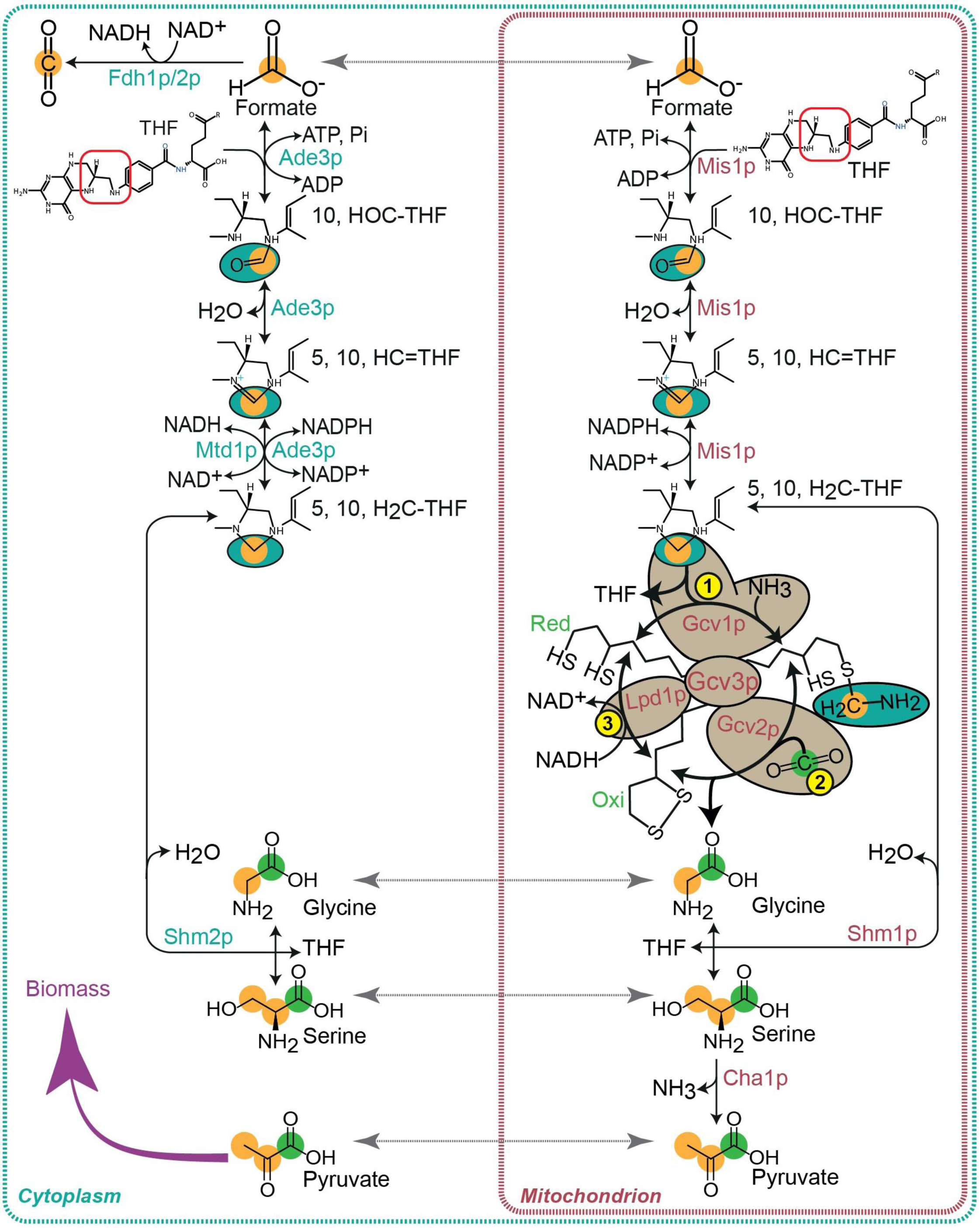
The Yeast RGP enzymes are distributed across the mitochondria and cytosol. The native enzymes that can form the RGP are found in different cellular compartments. Formate is ligated to mitochondrial THF by the trifunctional Mis1p enzyme to produce the mitochondrial pool of CHO-THF, CH-THF, and CH_2_-THF. In the cytoplasm, formate is ligated to cytoplasmic THF to produce the cytoplasmic pool of CHO-THF, CH-THF, and CH_2_-THF via the trifunctional Ade3p. The mitochondrial Glycine Synthase Complex (GSC) composed of Gcv1p, Gcv2p, Gcv3p, and Lpd1p mediates glycine synthesis by assimilating the CH_2_ group from CH_2_-THF, NH_3_, and CO_2_. Glycine synthesized in the mitochondria is converted to serine by assimilating another CH_2_-THF via Shm1p in the mitochondrial compartment and Shm2p in the cytoplasm. Serine is deaminated to pyruvate by mitochondrial Cha1p. **Mis1p**-mitochondrial trifunctional enzyme: formate-tetrahydrofolate ligase/ methenyltetrahydrofolate cyclohydrolase/ methylenetetrahydrofolate dehydrogenase; **Ade3p**-cytoplasmic trifunctional enzyme with identical functions of mitochondrial isozyme Mis1p; **Mtd1p:** cytoplasmic NAD+ dependent methylenetetrahydrofolate dehydrogenase that converts CH-THF to CH_2_-THF; **Gcv1p:** T subunit of the mitochondrial glycine decarboxylase complex (Aminomethyltransferase); **Gcv2p:** P subunit of the mitochondrial glycine decarboxylase complex (Glycine dehydrogenase); **Gcv3p:** H subunit of the mitochondrial glycine decarboxylase complex (carrier peptide hosting lipoamide); **Lpd1p:** Dihydrolipoamide dehydrogenase; **Shm1p:** mitochondrial serine hydroxymethyltransferase; **Shm2p:** cytoplasmic serine hydroxymethyltransferase; **Cha1p:** mitochondrial serine deaminase; **Fdh1p/2p:** formate dehydrogenases localized to cytoplasm.

For the second glycine-to-serine conversion module, the serine hydroxymethyltransferases, Shm1p and Shm2p, convert glycine to serine by combining glycine and the CH_2_-OH group from another formate-derived CH_2_-THF. As the C1-THF pools in mitochondria and cytoplasm are seperate, Mis1p provides CH_2_-THF molecules to facilitate mitochondrial glycine-to-serine conversion by Shm1p, whereas Ade3p supplies CH_2_-THF for cytosolic glycine-to-serine conversion by Shm2p. For serine synthesis, 2 ATP, 2 NADPH, and 1 NADH, along with two formate molecules, are required. The pyruvate synthesis module is driven by Cha1p, which catalyzes serine deamination to pyruvate in the mitochondria. Additionally, the endogenous formate dehydrogenases Fdh1p and Fdh2p allow for formate oxidation to CO_2_ and NADH, thereby providing reducing power and energy for the RGP. Based on the localization of Mis1p, Gcv1p, Gcv2p, Gcv3p, Lpd1p, Shm1p, and Cha1p, the entire assimilation pathway can operate in the mitochondria. While cytosolic Ade3p and Shm2p allow for CH_2_-THF synthesis from formate, and its assimilation into serine the full pathway cannot function in the cytoplasm. However, the energy module (Fdh1p and Fdh2p) is localized in the cytosol (**Figure 1**).

During yeast growth on glucose, the oxidation of serine and glycine for energy production is usually favored. Thus, the net flux of the reactions involving Shm1p, Shm2p, Gcv1-3p, Lpd1p, Mis1p, and Ade3p is towards releasing CO_2_ and formate. Formate can be shuttled between cytosol and mitochondria for C1-THF synthesis or oxidation. To facilitate formate and CO_2_ assimilation and to reverse the net flux toward glycine, serine, and pyruvate synthesis via the RGP, it is essential to provide elevated levels of CO_2_, in particular for reversal of the Gcv2p reaction^21^. Providing high formate and ammonium sulfate concentrations ((NH_4_)_2_SO_4_) might offer better kinetics for respectively CH_2_-THF synthesis by Mis1p and Ade3p, and for the amino-methyltransferase reaction via Gcv1p. In a previous study, we already demonstrated that the glycine synthesis module of the RGP can be engineered in *S. cerevisiae*, resulting in a strain capable of synthesizing glycine from formate in high CO_2_ conditions^25^. Here, we engineer a serine biosensor (ΔS) strain to implement the complete serine synthesis module of the RGP using a growth-coupled selection strategy^26^ combined with adaptive laboratory evolution (ALE), resulting in formate and CO_2_-dependent biosynthesis of glycine and serine. Further, we provide evidence for pyruvate synthesis, thereby demonstrating the carbon flux through the complete RGP in the *S. cerevisiae* for the first time.

## Results

### Engineering a serine biosensor strain

In order to establish glycine and serine synthesis from formate via the RGP, we applied a growth-coupled selection strategy employing an engineered serine biosensor (ΔS) strain. *S. cerevisiae* has three different glycine and serine biosynthesis routes, i.e., a predominant glycolytic serine biosynthesis pathway via 3-PGA during yeast growth on glucose^27,28^, a secondary serine biosynthesis pathway via glycine derived from threonine cleavage^29^, and a third pathway that is used to synthesize glycine from glyoxylate during growth on non-fermentable carbon sources like ethanol or acetate^30,31^. Glycine produced from the latter two pathways is converted to serine by condensing a hydroxymethyl group by Shm2p, the cytoplasmic serine-hydroxymethyltransferase (*SHMT*), and/or Shm1p, in the mitochondria (**Figure 2A**).

**Figure 2:**
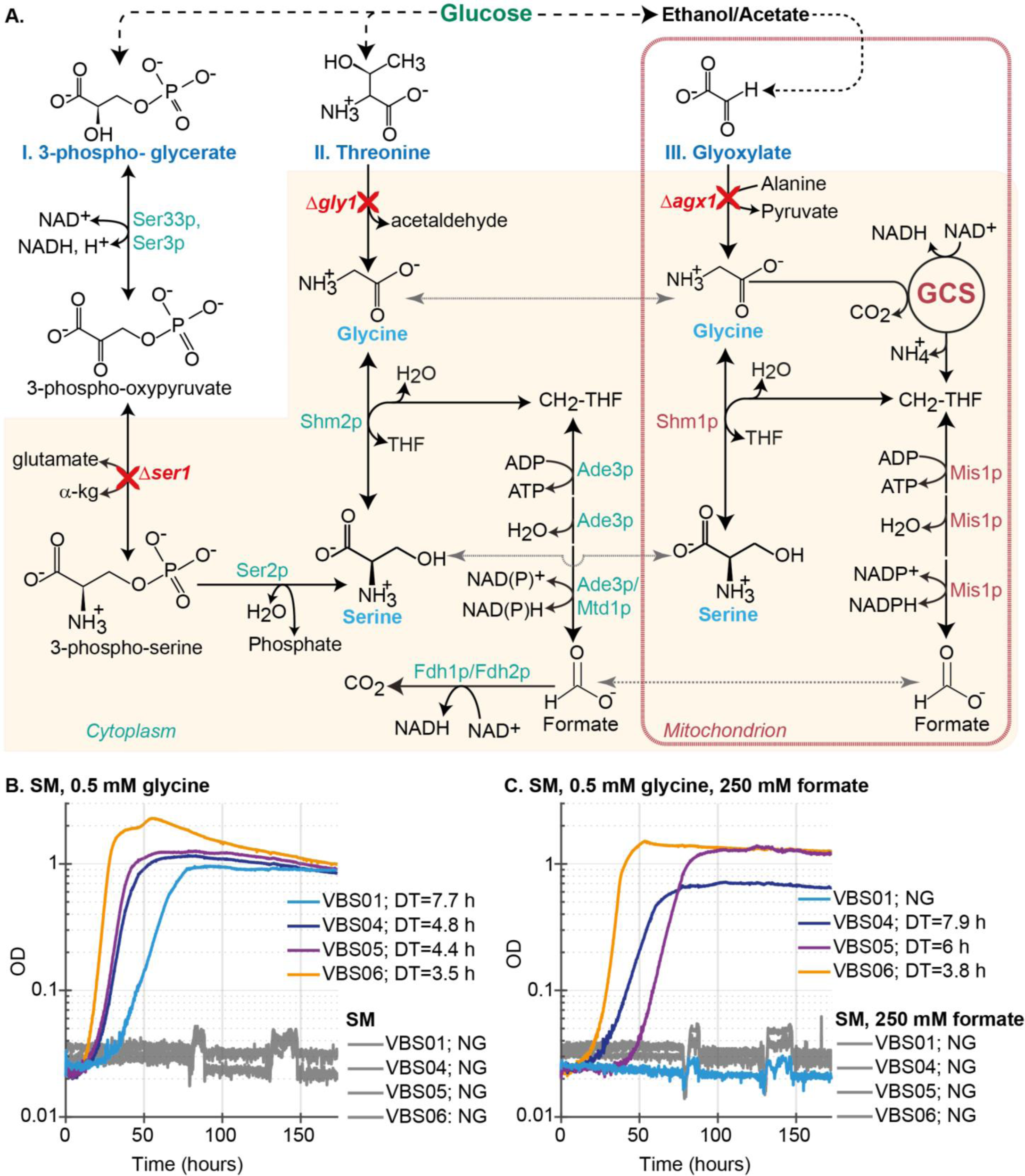
Construction of a serine biosensor strain and engineering glycine-to-serine conversion modules. A. Three different glycine and serine synthesis routes of yeast are shown. **I.** Serine synthesis via 3-PGA is the predominant pathway while yeast is growing on glucose. This pathway is disrupted by a *SER1* knockout. **II.** Threonine cleavage is a major pathway for glycine synthesis which is disrupted via *GLY1* knockout. **III.** *AGX1* is deleted to block the glycine flux via glyoxylate, which is active during growth on acetate or ethanol. Serine and glycine can be interconverted via Shm1p or Shm2 activity. Degradation of serine and glycine via Shm1p/Shm2p and GCS leads to CH_2_-THF synthesis, which is further oxidized to formate to release energy and reducing power via Ade3p and Mis1p. Formate is further oxidized to CO_2_ and NADH by Fdh1p/2p. **GCS:** Glycine Cleavage System; **Agx1p:** alanine-glyoxylate transaminase; **Gly1p:** Threonine aldolase; **Ser3p/Ser33p:** phosphoglycerate dehydrogenase; **Ser1p:** 3-phosphoserine aminotransferase; **Ser2p:** phosphoserine phosphatase; **B.** Growth of VBS01, VBS04, VBS05, and VBS06 strains in SM medium with or without glycine supplementation in a high CO_2_ atmosphere, in the absence of formate and **C.** in the presence of 250 mM formate. **SM:** 1x YNB, 40 mM (NH_4_)_2_SO_4_, 100 mM glucose; **YNB:** Yeast nitrogen base with ∼40 mM of (NH_4_)_2_SO_4_ **DT:** doubling time in hours. **NG:** No growth.

All three glycine and serine synthesis pathways were disrupted by creating complete knockouts of *AGX1* (Agx1p: Mitochondrial alanine:glyoxylate aminotransferase), *GLY1* (Gly1p: Cytosolic threonine aldolase), and *SER1* (Ser1p: Cytoplasmic 3-phosphoserine aminotransferase) from the genome to engineer a ΔS strain, i.e. *S. cerevisiae S288c Δagx1 Δgly1 Δser1* (VBS01) (**Figure 2A**). The combination of these deletions should result in a strain with a serine, glycine, and one-carbon network insulated from the central metabolism. As glycolytic flux cannot enter this isolated network, the strain cannot grow on glucose as the sole carbon source but requires an additional carbon source (glycine or serine) that can satisfy the auxotrophies for serine, glycine, and C1 metabolites.

We confirmed a strict auxotrophy for glycine and serine in the VBS01 strain by conducting growth assays in synthetic minimal (SM) medium with and without glycine or serine supplementation along with glucose as the primary carbon source. As expected, growth is only observed when at least one of these two amino acids is supplemented along with glucose, whereas the wild-type strain does not require supplementation of glycine or serine (**Figure 2B, Supplementary figure 1A**). If glycine is supplemented, the native glycine degradation through the GCS releases CH_2_-THF, which satisfies the cell’s need for C1 units. CH_2_-THF can be further condensed with another molecule of glycine and H_2_O by SHMTs (Shm1p or Shm2p) to yield serine in the mitochondria and/or in the cytosol. If serine is provided, the SHMTs activity is sufficient to produce glycine and CH_2_-THF to satisfy all three auxotrophies and allow growth (**Figure 2A & 2B, Supplementary figure 1A**). Our results confirm that VBS01 is a suitable ΔS strain with a simple growth/no growth readout, which should enable us to test the glycine and serine synthesis via a synthetic pathway. Further, since supplementation of glycine is sufficient to rescue growth, our observations suggest that the glycine to serine conversion via native Shm1p/Shm2p is unaffected by the introduced deletions. This is essential, since successful implementation of the RGP requires the conversion of glycine, eventually generated from formate and CO_2_, to serine.

However, once we cultured the biosensor strain in the conditions that are required for operating the RGP (SM medium with 250 mM formate as the secondary carbon source, 10% CO_2_ atmosphere), we observed that the glycine to serine conversion via the native Shm1p/Shm2p activity no longer supports growth of the VBS01 strain in the presence of formate and glycine (**Figure 2C, Supplementary figure 1B**). As opposed to this, serine supplementation is still sufficient for growth. This result indicates that the high formate concentration in the presence of glycine probably interferes with native SHMTs expression or activity directly or indirectly in the tested conditions, thereby preventing serine synthesis from glycine and formate – a core requirement for operating the RGP^32,33^. Thus, we overexpressed different SHMT submodules to test whether this enables growth via efficient glycine-to-serine conversion in the presence of formate and 10% CO_2_.

### Overexpressing SHMTs enables glycine to serine conversion under selective conditions

In order to enable glycine-to-serine conversion in the presence of 250 mM formate, we further engineered VBS01 and constructed three variants of the ΔS strains (**Supplementary table 1**) via genome integration of different SHMT overexpression (ox) cassettes, i.e., VBS04 with *SHM1*ox, VBS05 with *SHM2*ox, and VBS06 with *EcGlyA*ox, the SHMT from *E. coli*.

We conducted growth assays to test whether these strains can convert glycine-to-serine to support the growth with or without 250 mM formate addition. All three strains (VBS04-06) are able to grow with formate and glycine but otherwise retain their strict no-growth phenotype in the absence of glycine (**Figure 2B & 2C**). When supplied with glycine and formate, VBS04 and VBS05, overexpressing *SHM1* and *SHM2*, respectively, grow with doubling times of 7.9 h and 6 h, whereas VBS06, carrying the *EcGlyA* overexpression module, grows with a faster generation time of 3.8 h (**Figure 2B & 2C**) indicating efficient glycine-to-serine conversion by *Ec*GlyA. Nevertheless, all three strains, overexpressing one of the *SHMT* modules, can convert supplied glycine to serine in the presence of formate while incubated under a high CO_2_ atmosphere. This renders them (VBS04, VBS05, and VBS06) good candidates to further engineer the RGP.

These strains further require a glycine synthesis module to express the complete serine synthesis module of the RGP. Thus, we transformed pFM340 (**Supplementary table 2**), derived from pJGC3^25^ encoding the glycine synthesis module (*MIS1*, *GCV1-3)*, into these strains and generated VBS08 (*SHM1*ox), VBS09 (*SHM2*ox), and VBS10 (*EcGlyA*ox) strains (**Supplementary table 1**).

### Adaptive laboratory evolution enables enhanced glycine and serine biosynthesis from formate

We tested VBS08, VBS09, and VBS10 for their formate and CO_2_-dependent glycine and serine synthesis in SM media under a 10% CO_2_ atmosphere using glucose and formate as the primary and secondary carbon sources. As expected, none of the strains grew in SM media or SC-Gly-Ser (synthetic complete medium without glycine and serine) in the absence of formate, confirming that the auxotrophies are not affected by the expression of the glycine synthesis module (**Supplementary table 3**). To confirm that the SHMT modules are still functional in the strains carrying pFM340, we tested for growth in a medium supplemented with 0.5 mM glycine and 250 mM formate and observed growth in SM and SC-Ser medium for all three strains. Finally, to test the activity of the glycine synthesis module and the SHMTs, we inoculated these strains in medium supplemented with formate as the secondary carbon source. However, only one of the three strains, VBS10, grew to a very low OD under the conditions tested (**Supplementary figure 2A**). This growth was only observed in the SC-Gly-Ser medium supplied with formate and glucose but not in the SM medium with formate and glucose (**Supplementary table 3**). This result suggests at least some glycine and serine biosynthesis from formate.

Considering this as the first indication of glycine and serine synthesis from formate, we designated this VBS10 culture as Evolution round 01 (Ev01) and proceed through Adaptive Laboratory Evolution (ALE) to establish stable formate and high CO_2_ dependent growth in minimal media (**Figure 3A & 3B**). We diluted cells from Ev01 into fresh media – again in SC-Gly-Ser and SM media, both supplemented with formate as the secondary carbon source. Growth was observed in Ev02 only after a prolonged incubation of ∼30 days in SC-Gly-Ser medium provided with formate and glucose, which was used as seed inoculum in Ev03 (**Figure 3A**). Here, growth was observed in the SM medium supplied with glucose and formate, but not in the absence of formate (**Figure 3A & 3C**). At this stage of ALE, the growth rate was slow, with a doubling time of ∼80 h, in SM medium. Thus, we continued the ALE process by passaging into the next cycles in a fresh SM medium supplied with formate as a secondary carbon source. Growth only occurred in medium supplemented with formate and glucose under a high CO_2_ atmosphere. This indicates that the observed growth is due to glycine and serine biosynthesis from formate and CO_2_ via the RGP (**Supplementary figure 2B**). In order to improve the fitness of the strain, we continued the evolution by diluting the grown cells from the late exponential phase to the subsequent Ev culture. Over the course of 21 cycles of ALE, we observed a decrease in the apparent doubling time of the VBS10 strain from ∼80 h in Ev03 to as low as ∼7 h (**Figure 3C**).

**Figure 3:**
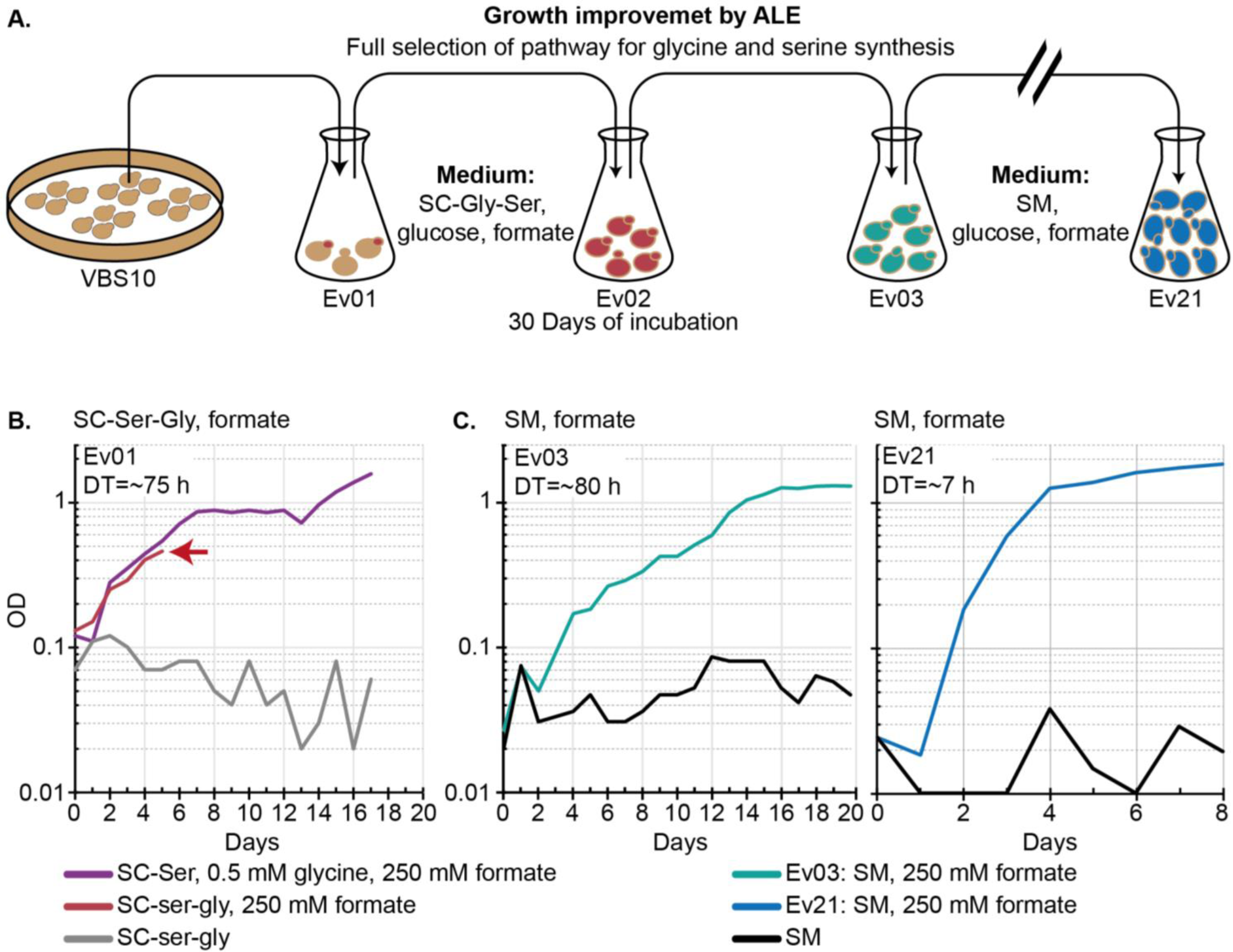
Adaptive Laboratory Evolution of VBS10 in semi-rich medium to minimal medium. **A.** Original VBS10 transformants were pooled and inoculated in SC-Gly-Ser medium supplemented with formate (Ev01). The cells from Ev01 were sub-cultured in the same medium in Ev02, and growth was observed with prolonged incubation. VBS10_Ev02 grown cells were then inoculated to SM medium supplemented with formate (Ev03). The population from each Ev was used as seed for subsequent rounds up to Ev21 to improve growth. **B.** Growth of VBS10_Ev01 in SC-Gly-Ser medium supplemented with/without formate and glycine+formate. **C.** Growth of VBS10 in Ev03 and Ev21 in SM medium supplemented with/without formate. Cultures were always incubated under 10% CO_2_ atmosphere while testing for glycine and serine synthesis from formate.; **DT:** doubling time in hours; All growth curves are average values of duplicates of each selection condition across the ALE.

For further analysis, we isolated individual clones from ALE cycles of Ev03, Ev06, Ev12, Ev16, and Ev21. Microtiter plate-based growth assays were performed for 10 clones of each ALE cycle. The mean growth rate for isolated clones between Ev03-Ev21 cycles significantly increased from 0.009 h^-1^ to 0.085 h^-1^ (average doubling time decreased from ∼75 h to ∼8 h) (**Figure 4**). A significant growth improvement was observed in each selected evolution round compared to its selected predecessor round until Ev16, with the highest growth improvement between Ev03 and Ev06. Growth improvement between Ev16 and Ev21 is insignificant, but the variance of growth rates among the Ev21 and Ev16 clones is decreased from 1×10^-4^ to 1×10^-5^. The decrease in the variance between individual clones indicates that either a single strain slowly took over the culture or multiple strains converged against a similar growth rate. In all experiments, we never observed growth without formate or CO_2_, indicating that formate assimilated via the RGP is the source of glycine and serine in the evolved strains (**Figure 4**).

**Figure 4:**
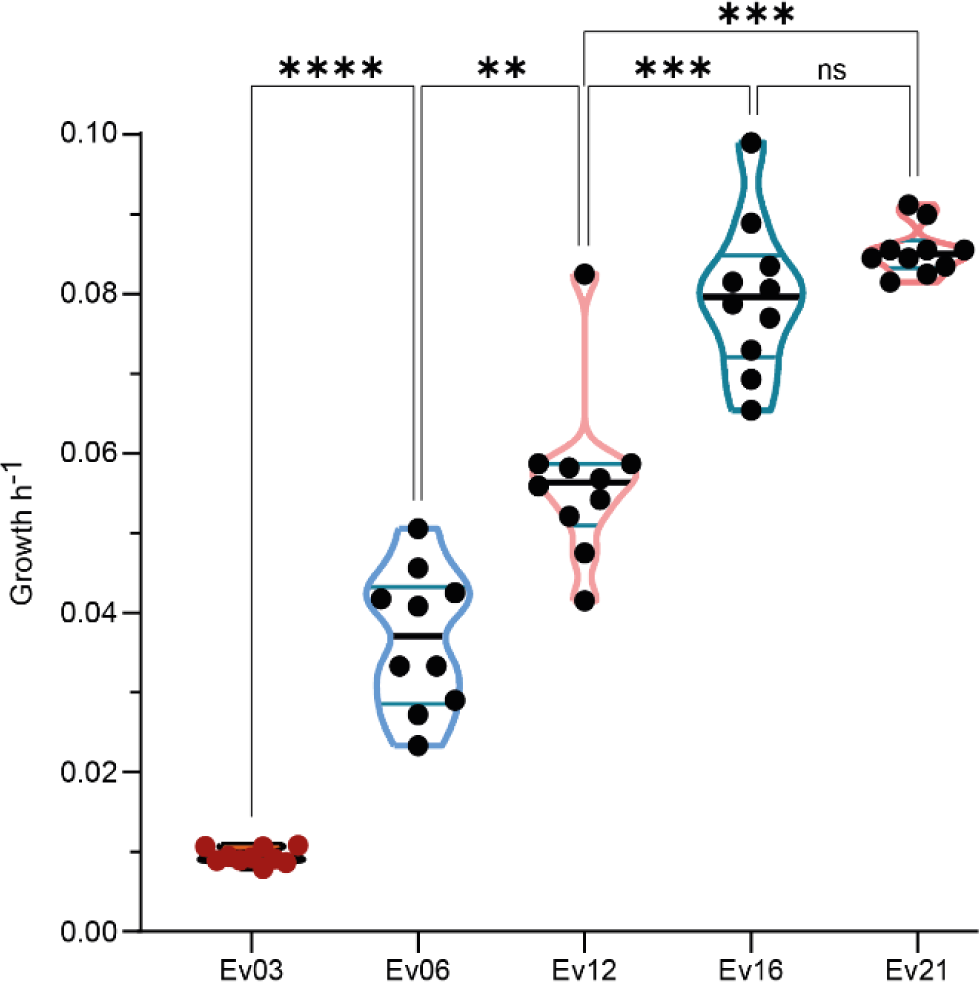
Growth improvement of VBS10 strains during ALE. Growth rates of multiple clones isolated from evolved VBS10 populations Ev03, Ev06, Ev12, Ev16, and Ev21 were assessed. Mean difference of growth rates between Ev03 and Ev21 is 0.0752 h^-1^. One-way ANOVA was performed to assess the significance of the growth rate improvements during ALE and the p-value was calculated. The significance of the growth improvement is indicated with stars based on p -values. p=<0.0001, is indicated as ****, *** indicates the p=0.0002-0.0005, p=0.0014 is indicated as **, and p=0.0338 as *, and no significance was indicated as ‘ns - p=>0.05’.

### Validation of glycine and serine biosynthesis from formate by isotope labeling

To investigate if the observed growth is indeed due to glycine and serine synthesis from formate, we conducted ^13^C-labeling experiments with evolved VBS10 and WT strains. We grew both strains with either ^12^C or ^13^C formate in SM medium with ^12^C glucose as a primary carbon source in a 10% ^12^CO_2_ or ^13^CO_2_ atmosphere. The biomass of the WT and evolved VBS10 was analyzed for ^13^C-carbon in proteinogenic glycine, serine, and alanine using LC-MS. As shown in **Figure 5** our results match the expected labeling pattern for glycine and serine biosynthesis from formate and CO_2_ via the RGP.

**Figure 5:**
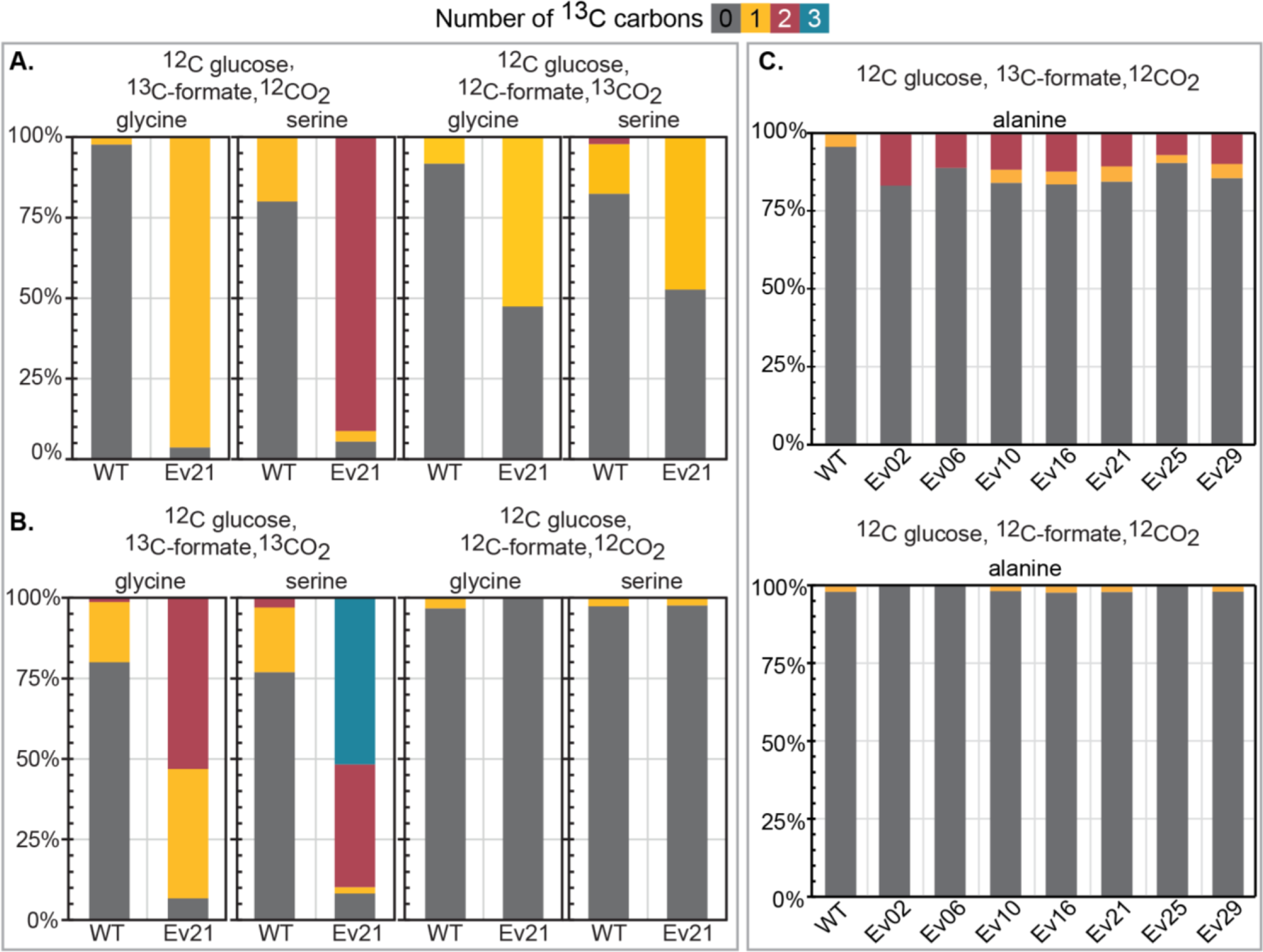
^13^C-based carbon tracing experiments validate the RGP activity in the evolved VBS10 strains. The fractions of labeled proteinogenic glycine and serine from the biomass of evolved VBS10 and wild-type strains **A.** Cells grown with ^12^C-glucose + ^13^C-formate + ^12^CO_2_, and ^12^C-glucose + ^12^C-formate + ^13^CO_2_. **B.** Cells grown with ^12^C-glucose + ^13^C-formate + ^13^CO_2_, and ^12^C-glucose + ^12^C-formate + ^12^CO_2_. When VBS10 is fed with ^13^C formate, the near complete ^13^C_1_-glycine, and ^13^C_2_-serine labelling confirms formate assimilation into glycine and serine. When ^13^CO_2_/^13^CO_2_+^13^C-formate is fed, only around 50% of glycine and serine carry labels in all carbon atoms. The rest of the incompletely labeled glycine and serine is due to the competitive assimilation of glycolytic ^12^CO_2_ generated in mitochondria and ^13^CO_2_ from the incubation atmosphere. **C.** Fractions of ^13^C_2_-alanine indicate the transfer of two carbons (^13^C_2_) from ^13^C-formate-derived ^13^C_2_-serine via deamination to ^13^C_2_-pyruvate and transamination to ^13^C_2_-alanine, validating pyruvate synthesis from formate via RGP (**Supplementary** figure 3). No ^13^C-labeling is observed when strains are fed with only^12^C-carbon sources.

When ^13^C formate is fed, >95 % of all glycine is once-labeled in all evolved strains. When ^13^CO_2_ is fed, around 50 % of the glycine is single-labeled, while the remainder is unlabeled. Consistently, when ^13^CO_2_ and ^13^C-formate are fed, around 50 % of glycine is dual-labeled (**Figure 5A & 5B**). Together, these results prove the activity of the RGP core module, resulting in glycine biosynthesis from formate. Detailed visualization of ^13^C carbon fate is provided in the **Supplementary figure S4**. The observation that the ^13^CO_2_ is only partially incorporated in glycine is very likely due to the high amount of ^12^CO_2_ produced from ^12^C glucose in the mitochondria via the TCA and the pyruvate dehydrogenase.

Nearly all serine (>90 %) in the biomass of the evolved VBS10 strains is dual-labeled when supplied with ^13^C-formate. This demonstrates the formate-derived glycine conversion to serine via the assimilation of ^13^CH_2_-THF derived from formate, validating the activity of the complete serine synthesis module of the RGP. When either ^13^CO_2_ or ^13^CO_2_ and ^13^C-formate were supplied, the labeling pattern of serine again showed partial incorporation of glycolytic ^12^CO_2_ resulting in ∼50% of ^13^C_2_-serine or ^13^C_3_-serine, respectively. Per approximately every two molecules of glycine/serine synthesis, one molecule of the atmospheric CO_2_ (from the incubation setup) and one CO_2_ molecule coming from oxidized glucose is assimilated (**Figure 5A & 5B**).

As expected, for both glycine and serine, the wild type control strain does not carry labeling comparable to the strain operating the RGP when fed with ^13^CO_2_ or ^13^CO_2_ and ^13^C-formate (**Supplementary figure 3**). The ^13^C_1_-serine fraction in the wild type, indicates that part of ^13^CH_2_-THF is derived from the supplied ^13^C-formate and is assimilated to make ^13^C_1_-serine from non-labelled glycine (coming from threonine or glyoxylate). In the case of ^13^CO_2_, carbon fixing anaplerosis reactions might result in a fraction of ^13^C_1_-glycine via threonine cleavage catalyzed by Gly1p. Considering these observations, our results confirm that serine and glycine are exclusively produced from formate with a partial CO_2_ assimilation from the incubation atmosphere via the RGP in the evolved ΔS strains. As expected, we have observed the nearly complete supply of C1 units for histidine and methionine synthesis from formate in the ΔS strains demonstrating the isolation of the C1-network, as opposed to the wild type (**Supplementary figure 4**). Thus, all the C1-units, glycine and serine are synthesized via the RGP in the evolved VBS10 strains.

Additionally, we observed around ∼10%-15% of proteinogenic ^13^C_2_ labelled alanine in the evolved VBS10 strains when fed with ^13^C-formate (**Figure 5C**). As alanine is synthesized from pyruvate these results indicate that also a fraction of the cellular pyruvate is derived from formate, even though there is no growth-coupled selection pressure for pyruvate synthesis via the RGP when growing the strains on formate and glucose. This means that formate-derived ^13^C_2_-serine is deaminated to ^13^C_2_-pyruvate via native Cha1p and is further trans-aminated via native Alt1p, leading to 10%-15% ^13^C_2_-alanine (**Supplementary figure 3**) across different isolates form ALE experiments, although there is no selection pressure for pyruvate synthesis (**Figure 5C**). This suggests that all three modules of the RGP are active in the evolved VBS10 and is the first indication that the full RGP operates in the yeast. Thus, it was tempting to test if the strain could grow with formate as the sole carbon and energy source. However, we could not detect growth when the strain was inoculated in SM with 250 mM formate as the sole carbon source. Thus, we speculated that the amount of pyruvate generated from formate is insufficient to support full formatotrophic growth.

To address this potential limitation, we applied short-term ALE to enhance the flux towards pyruvate. For this, we grew the cells over eight ALE cycles with decreasing glucose concentrations in SM medium supplied with 100 or 250 mM formate. The final OD of these cultures was reduced with decreasing glucose concentrations - irrespective of the formate concentration, suggesting that glucose is the limiting compound at this stage (**Supplementary figure 5**). However, carbon tracing experiments with isolates from Ev25 and Ev29 fed with ^13^C-formate still show a ∼10% fraction of ^13^C_2_-alanine, as observed for previous isolates. This again shows partial conversion to pyruvate from formate, but apparently without improvements in the serine to pyruvate flux (**Figure 5C**). We continued the ALE in 2mM glucose up to Ev50, but growth was not improved, irrespective of the formate availability. In addition, elongated cell growth phenotype on SM media plates under 2 mM glucose and 100 mM formate indicates energy deficiency. These results suggest that there is a lack of or insufficient formate dehydrogenase activity to produce NADH. Hence, VBS10 may be unable to use formate to generate enough energy and reducing power, leading to inadequate serine availability for pyruvate synthesis. Summarizing, ALE in glucose-limiting conditions did not improve the pathway flux, thus, a further engineering and evolution approach is essential.

### Substrate dependency of the RGP in the evolved VBS10

The RGP is operated with three substrates, i.e., formate and CO_2_, as carbon sources and ammonium sulphate ((NH_4_)_2_SO_4_) as the nitrogen source. One molecule of all three substrates is assimilated to produce one glycine molecule, and another formate molecule (ligated to THF and reduced to CH_2_-THF) is required for the glycine conversion to serine. When serine is deaminated to pyruvate, released ammonia molecules can be recycled in the next round of glycine synthesis. To understand which of the three substrates is important for the RGP-dependent growth of evolved strains, we subjected VBS10_Ev21 strains to a series of growth experiments with concentration gradients of formate, ammonium sulfate, and varying percentages of elevated CO_2_ in the incubation atmosphere.

The VBS10_Ev21 strains grow on formate concentrations as low as 10 mM and up to 750 mM of formate with average growth rates ranging from ∼0.03 h^-1^ to ∼0.1 h^-1^. Best growth rates are observed between 50 mM to 125 mM, which might provide optimum kinetics for the RGP operation with minimal formate toxicity. However, below 25 mM and above 300 mM, a sharp drop in the growth rates can be observed (**Figure 6A**). While the latter is most likely due to formate toxicity, the first can be interpreted as a limited driving force for RGP-dependent growth. Following a similar trend, the biomass yield based on final OD_600_ observed at formate concentrations between 50 mM and 125 mM, is highest (**Supplementary figure 6A**).

**Figure 6:**
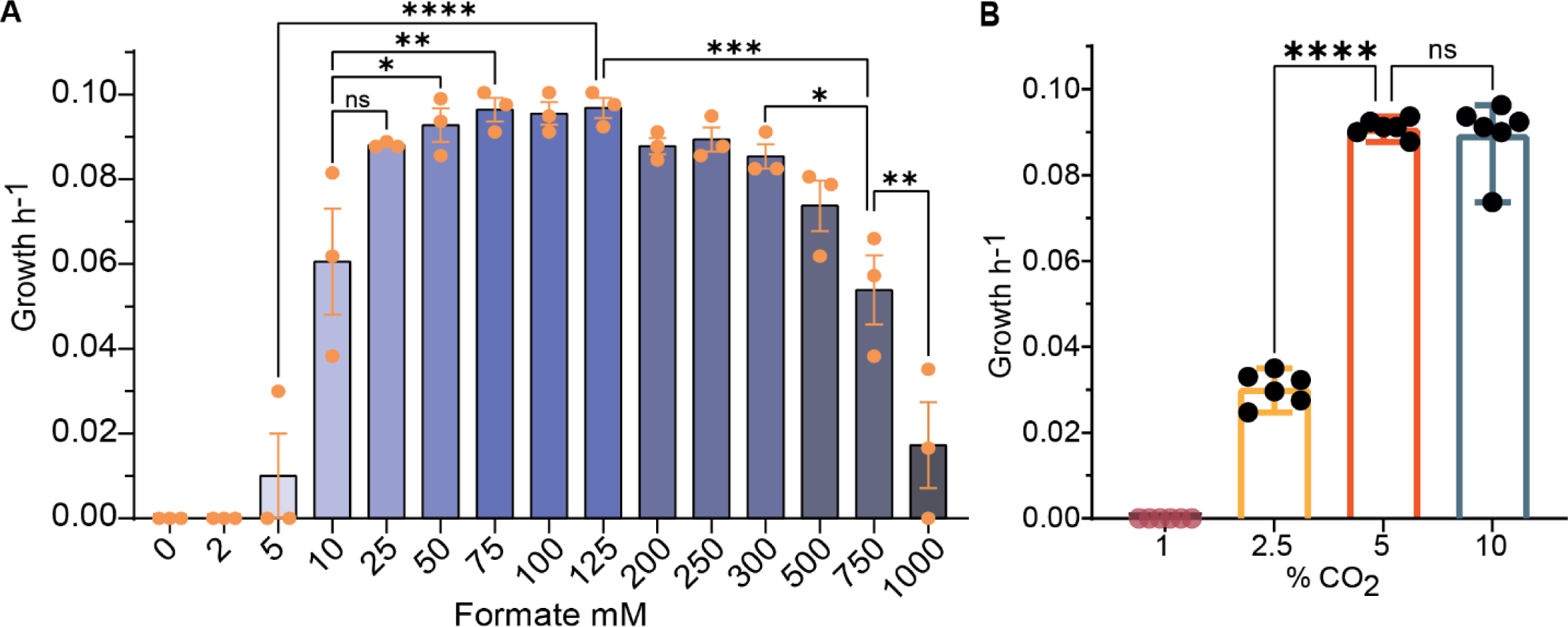
Effect of formate and CO_2_ on RGP-dependent growth of the evolved VBS10 strains. **A.** Growth of VBS10_Ev21 strains in SM medium with various formate concentrations as the secondary carbon source, incubated under high CO_2_ atmosphere. **B.** Growth of VBS10_Ev21 when incubated in different atmospheres from 10% to 1% CO_2_. The Growth rate of VBS10_Ev21 sharply dropped when the CO_2_ percentage in the incubation atmosphere was reduced from 5% CO_2_ to 2.5% CO_2_. All growth experiments were conducted in an SM medium supplied with glucose as the primary and formate as the secondary carbon sources.

The VBS10 strains cannot grow via the RGP in ambient CO_2_ conditions, an elevated CO_2_ atmosphere is necessary for RGP operation. To determine the minimal CO_2_ concentration essential for RGP-dependent growth of the evolved VBS10 strains, we assessed VBS10 growth with atmospheric CO_2_ concentrations between 1 % and 10 %. VBS10 grows with all CO_2_ levels tested, except for 1 % CO_2_ (**Supplementary figure 6B**). A significant decrease in the growth rate is observed with 2.5 % CO_2_, while 5 % CO_2_ is high enough to achieve the maximal growth rate (**Figure 6B**). Probably, 5% CO_2_ can provide sufficient glycine synthase activity via the rGCS. Nonetheless, VBS10 still grows with a reduced growth rate when CO_2_ is reduced to 2.5%, indicating the reversibility of the GCS at 2.5% CO_2_ for glycine synthesis in yeast. This development can be a starting point for future evolution experiments aiming to improve growth in low CO_2_ conditions employing different engineering and evolution strategies.

Varying the (NH_4_)_2_SO_4_ concentration in the 20 mM – 100 mM range (0.5 x – 3 x of the standard concentration in SM medium) did not influence the growth rate noticeably, however, 10 mM (0.25x) (NH_4_)_2_SO_4_ significantly diminished the growth rate in Ev21 compared to WT in the same condition (**Supplementary figure 6C & 6D**). Thus, lowering the (NH_4_)_2_SO_4_ below 20 mM slows down RGP-dependent growth, possibly due to the reduced aminotransferase reaction of GSC, lowering the glycine and serine synthesis rate. These results align with previous observations which show that a lower concentration of (NH_4_)_2_SO_4_ may affect the RGP-dependent growth of *D. desulfuricans*^20^.

### Identification of potential key mutations for formatotrophic growth

We performed whole genome sequencing for evolved VBS10 strains with the best growth rates, isolated from selected ALE rounds. Using the unevolved parental strain as a control, we identified several mutations acquired during the ALE process by sequencing isolates from different steps of the evolution (**Supplementary table 4**). A 36 amino acid deletion in the glutamate dehydrogenase 1 (*GDH1::Δ108bp)* mutation was presented in all sequenced isolates during the evolution. Several other mutations in coding regions and intergenic regions accumulated later during ALE (**Supplementary table 4**). We speculate that the amino acid deletions in glutamate dehydrogenase 1 (*Gdh1p::Δ36aa* mutation) and amino acid substitutions in isocitrate dehydrogenase 1 (Idh1p-Pro275Gln), within the AMP binding domain (**Supplementary figure 7A-7D**), and the major ADP/ATP carrier protein on the mitochondrial inner membrane, *PET9* (Pet9p-Ile275Thr)^34,35^, might play a role in the re-distribution of energy and reducing power, promoting RGP operation. Among these three mutations, the *gdh1* knockout mutation was identified right from the Ev01/02, and both other mutations were identified from Ev06. The mutations Utp10-Met1Ile (in a nucleolar protein involved in pre-18s-rRNA processing), identified from Ev06, Sui2p-Arg88Leu (in α-subunit of the translation initiation factor eIF2), and *ASH1::Δ3pb* (in GATA-like transcription factor), the latter two both identified in the Ev21, are potential regulatory mutations that are speculated to contribute to the fitness of evolved VBS10 strains. Further, intergenic mutations with potential effects on the expression of their adjacent genes were found interesting for glutamate cysteine ligase (Gsh1p) and Met6p (Cobalamin-independent methionine synthase), among others. These mutations maybe of interest of interest because of their early evolution appearance and their functional connection to glutamate and one-carbon metabolism, respectively.

### Reverse engineering of *gdh1::Δ108bp* deletion mutation

The 108bp deletion in the CDS of *GDH1* between 222 and 330 nucleotides identified throughout clones from all evolution rounds translates to a truncated glutamate dehydrogenase 1, Gdh1p::*Δ*36aa. Its early appearance in all the clones from the VBS10_Ev01/02 suggests a significant role in enabling the RGP. Based on the Gdh1p domain structure, it is likely that the resulting truncated protein is non-catalytic, as substrate and cofactor binding regions are lost. The cytoplasmic Gdh1p, drives glutamate synthesis from α-ketoglutarate (α-KG) and ammonium, using NADPH as a cofactor. Nevertheless, Gdh1p::*Δ*36aa does not have to be detrimental in yeast as its cytoplasmic paralog, Gdh3p, catalyzes the same reaction at a lower rate. The latter is known to be induced by ethanol or other non-fermentable carbon sources^36,37^.

To investigate if this mutation can facilitate glycine and serine biosynthesis via the RGP, we introduced the mutation in unevolved VBS10 using CRISPR-Cas9. Unevolved VBS10 and VBS10 *gdh1::Δ108* (VBS18) strains were inoculated in SC-Gly-Ser medium with or without formate supplementation as the secondary carbon source. As expected, both strains did not show immediate growth. However, we noticed that all four clones of VBS18 reached a high OD_600_ after 25 days of incubation under high CO_2_ atmosphere, while VBS10 did not grow even in 40 days, until the end of the experiment (**Supplementary figure 8**). This suggests that the *gdh1::Δ108* plays a direct or indirect role in enabling growth-sufficing flux through the RGP in VBS18. However, the required longer incubation, over 25 days is expected, as growth in Ev02 of the ALE process also needed a longer incubation period and the doubling time was about 3 days (**Figure 3A & 3B**). It can be speculated that a loss of Gdh1p activity provides an additional NADPH pool available for the NADPH-consuming RGP, which might result in enhanced CH_2_-THF synthesis (**Figure 1**), thus enabling sufficient biosynthesis of glycine and serine to overcome auxotrophies.

To check if the *gdh1::Δ108pb* leads to increased CH_2_-THF synthesis, we introduced the same knockout in the WT background, and performed comparative formate assimilation between S288c WT and S288c *gdh1::Δ108bp* (VBS19; **Supplementary table 1)**. In the C1 network, histidine derives carbon from CHO-THF via purine metabolism, CH_3_-THF produced from CH_2_-THF provides carbon for methionine, and SHMTs assimilate carbon directly from CH_2_-THF into serine (**Figure 7A**)^38–40^. If there is any effect of *gdh1* inactivation on CH_2_-THF synthesis from formate, it should reflect in the ^13^C-labeling of histidine, methionine, and serine associated with the C1 network in yeast. To check this, we performed ^13^C-formate labeling experiments to compare the formate-derived CH_2_-THF and CHO-THF assimilation between both strains, S288c WT and VBS19.

**Figure 7:**
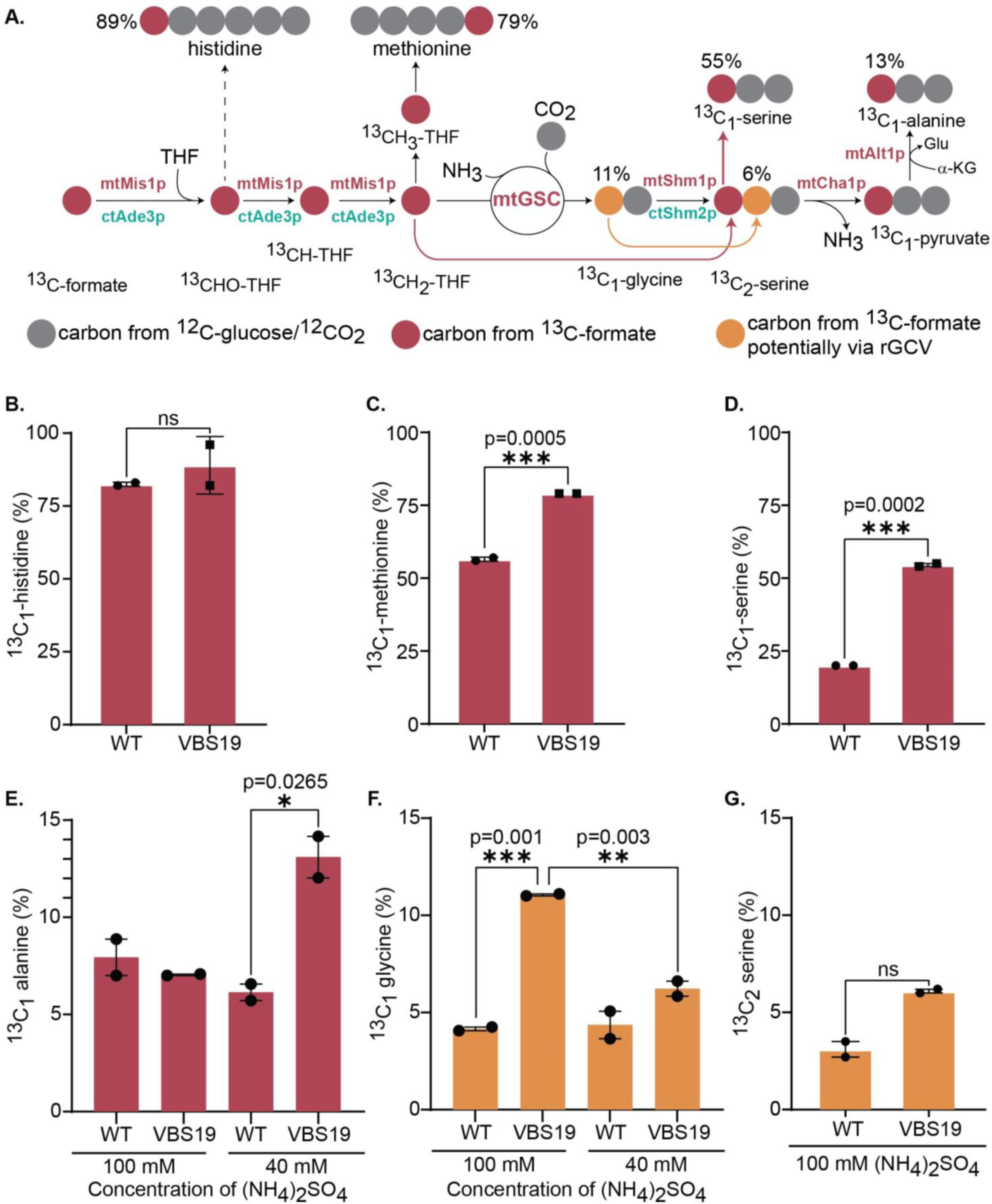
Comparative formate assimilation between VBS19 and S288c WT control. **A.** Formate assimilation in the C1-network of wildtype *S. cerevisiae* leading to carbon transfer form of ^13^C-formate to histidine, methionine, serine, alanine (pyruvate), and glycine. Percentage of **B.** ^13^C_1_-histidine, **C.** ^13^C_1_-methionine, and **D.** ^13^C_1_-serine in the VBS19 strains and the WT control. Percentage of **E.** ^13^C_1_-alanine via formate derived ^13^C_1_-serine, **F.** ^13^C_1_-glycine potentially via reversal activity of GCS **G.** ^13^C_2_-serine from ^13^C_1_-glycine in the VBS19 strain against WT control.; ***Δgdh1*** *= gdh1::Δ108bp***; WT** = S288c WT**; VBS19** = *S288c gdh1::Δ108bp*. Mt-mitochondrial, ct-cytosolic.

For this, we fed both strains with 250 mM ^13^C-formate, along with low ^12^C-glucose (5 mM), to minimize the interference of glucose-derived C1 assimilation because the one-carbon network is not insulated from the glycolytic flux in both these strains, unlike an ΔS strain (**Figure 2A**), in which all the C1-units are produced from externally supplied formate (**Supplementary figure 4**). The proteinogenic amino acids from the biomass of the S288c WT and VBS19 were analyzed for the differences in the ^13^C-labeling of histidine, methionine, and serine. Significantly higher percentages of ^13^C_1_-methionine and ^13^C_1_-serine were found in VBS19. In contrast, the difference in the percentage of ^13^C_1_-histidine is insignificant (**Figure 7B, C & D; Supplementary figure 9**. These results suggest increased synthesis of H_2_C-THF units derived from formate in the VBS19 strain compared to the S288c WT control. CH_2_-THF synthesis from formate is NADPH-dependent and hence is boosted by the *GDH1* deletion. This hypothetical role of the *GDH1* deletion in NADPH-dependent C1-biosynthesis is further supported by the fact that CHO-THF synthesis from formate (as reflected in histidine) is not increased in the *gdh1::Δ108bp* mutant. CHO-THF biosynthesis from formate does not involve NADPH consumption and hence is likely not influenced by the mutation. Our data confirm that the *gdh1::Δ108bp* stimulates NADPH-dependent C1-biosynthesis, likely by increasing the available NADPH pool in the cell and hence this mutation supports activity of the RGP (**Figure 1 & Figure 7A**).

About 12% of ^13^C_1_-alanine is observed in VBS19, which is significantly more than in the WT control. These differences were observed when 40 mM (NH_4_)_2_SO_4_ (∼1x) was provided as the nitrogen source (**Figure 7E**). Any fraction of ^13^C_1_-alanine can be attributed to the ^13^C_1_-serine that is deaminated to ^13^C_1_-pyruvate, which is further trans-aminated to ^13^C_1_-alanine, ultimately assimilating ^13^C_1_ from formate-derived ^13^CH_2_-THF. When 100 mM (NH_4_)_2_SO_4_ (∼2.5x) is supplemented in the medium, the difference in ^13^C_1_-alanine fractions is insignificant between VBS19 and WT, which is most likely due to the inhibition of the serine deamination due to high ammonium availability. In a contrasting phenomenon, significantly more ^13^C_1_-glycine was observed in VBS19 compared to the WT control when 2.5x (NH_4_)_2_SO_4_ is supplemented in the medium but not otherwise (**Figure 7F**). This probably translated to an insignificant rise in the ^13^C_2_-serine fraction in the VBS19 strain, only when a high nitrogen source was provided (**Figure 7G**). Nevertheless, this contrary effect of high NH_3_ on glycine and pyruvate synthesis can be minimized or nullified in an evolved strain operating the full RGP where the net NH_3_ assimilation is zero; thus, RGP-dependent growth might not require high (NH_4_)_2_SO_4_ once the full growth is achieved. The increased ^13^C_1_-glycine fraction with 2.5x (NH_4_)_2_SO_4_ indicates improved glycine synthase activity which is reported to be influenced by (NH_4_)_2_SO ^20^. Though a significant Fdh1p/2p activity was not observed in the evolved VBS10 (see above), minimal formate oxidation to CO_2_ cannot be ruled out. Thus, anaplerotic assimilation of the formate-derived CO_2_ might lead to ^13^C_1_-glycine via ^13^C_1_-threonine cleavage. ^13^C_1_-glycine flux via latent rGCS activity and possible ^13^C_1_-threonine cleavage are not insulated and thus cannot be distinguished. Nevertheless, the ^13^C_1_-glycine flux is only observed when high (NH_4_)_2_SO_4_ is provided. Since the RGP-dependent growth is influenced by (NH_4_)_2_SO_4_ in bacteria^20^, this flux is likely via rGCS without over-expression of the RGP in VBS19.

In the VBS10 strain, glycine, serine, and one-carbon metabolism are insulated as it is a ΔS strain carrying over-expression cassettes of the complete serine synthesis module of the RGP. Hence, all the C1-units are generated from the formate in the VBS10 (**Supplementary figure 4**). Our finding suggests that Gdh1p inactivation led to increased CH_2_-THF flux, which is one of the critical requirements for rGCS along with CO_2_. This, combined with prolonged incubation might lead to the redistribution of C1-units and NADPH via metabolic shunts between the cytoplasm and mitochondria. Eventually, leading to the GSC activity of mitochondrial rGCS, resulting in glycine, serine, and pyruvate synthesis from formate in the high CO_2_ atmosphere^40,41^.

## Discussion

This study, for the first time, demonstrates activity of the complete RGP in yeast, transferring carbon from formate to pyruvate. We successfully developed a *S. cerevisiae* S288c based strain, capable of producing all glycine and serine from formate and CO_2_, along with a fraction of pyruvate through the RGP. Using a growth-coupled biosensor strain of *S. cerevisiae,* we combined rational engineering and ALE. We first constructed a base ΔS (VBS01) strain and then implemented functional glycine-to-serine submodules in the RGP-specific conditions. ALE with three different ΔS strains carrying a complete serine synthesis module enabled the RGP operation in VBS10, but not in VBS08 and VBS09. Our carbon tracing experiments validate the carbon transfer from formate to pyruvate in the evolved VBS10 strain.

As shown in Figure 8, the evolved VBS10 strain currently operates an overexpressed glycine synthesis module in the mitochondria and the glycine-to-serine conversion submodule in the cytoplasm. Ade3p serves to supply the cytosolic CH_2_-THF pool for glycine-to-serine conversion. Thus, it is likely that the majority of the RGP fluxes might be as follows: glycine produced in the mitochondria is transported to the cytosol for serine synthesis. Serine is then imported to mitochondria for pyruvate synthesis via Cha1p^42^ and transamination to alanine via Alt1p. However, basal levels of mitochondrial glycine-to-serine conversion might still present via Shm1p^33^ which could also contribute to pyruvate synthesis. Combining all together this strain operates a complete serine synthesis module spanning across mitochondria and cytoplasm. Besides the consistent evidence for pyruvate synthesis in the form of alanine across different ALE stages of VBS10, the pyruvate flux seems insufficient to support total biomass and energy regeneration, and glucose-limiting ALE did not improve it further. We speculate that the lack of or inadequate formate dehydrogenase activity is a primary limitation. Though *S. cerevisiae S288c* has *FDH1* and truncated *FDH2*, apparent energy support for RGP was not evident in the tested conditions. However, basal activity of the Fdh may still be present, which may not generate enough NADH to support glycine synthesis in the mitochondria. Thus, it is necessary to establish an efficient Fdh activity e.g., from a methylotrophic yeast like *Candida boidinii* and target it to mitochondria for energy generation^43^. Overexpression of *CHA1* and *EcSdaA* to establish efficient serine deamination for pyruvate synthesis both in mitochondrial and cytosol and further evolution will be required to improve the flux towards pyruvate (**Figure 8**). This overexpression could overcome regulatory constraints of *CHA1* expression by *CHA4*, the transcriptional activator of yeast serine deaminase, in response to hydroxy-amino acids, serine, and threonine^42^.

**Figure 8:**
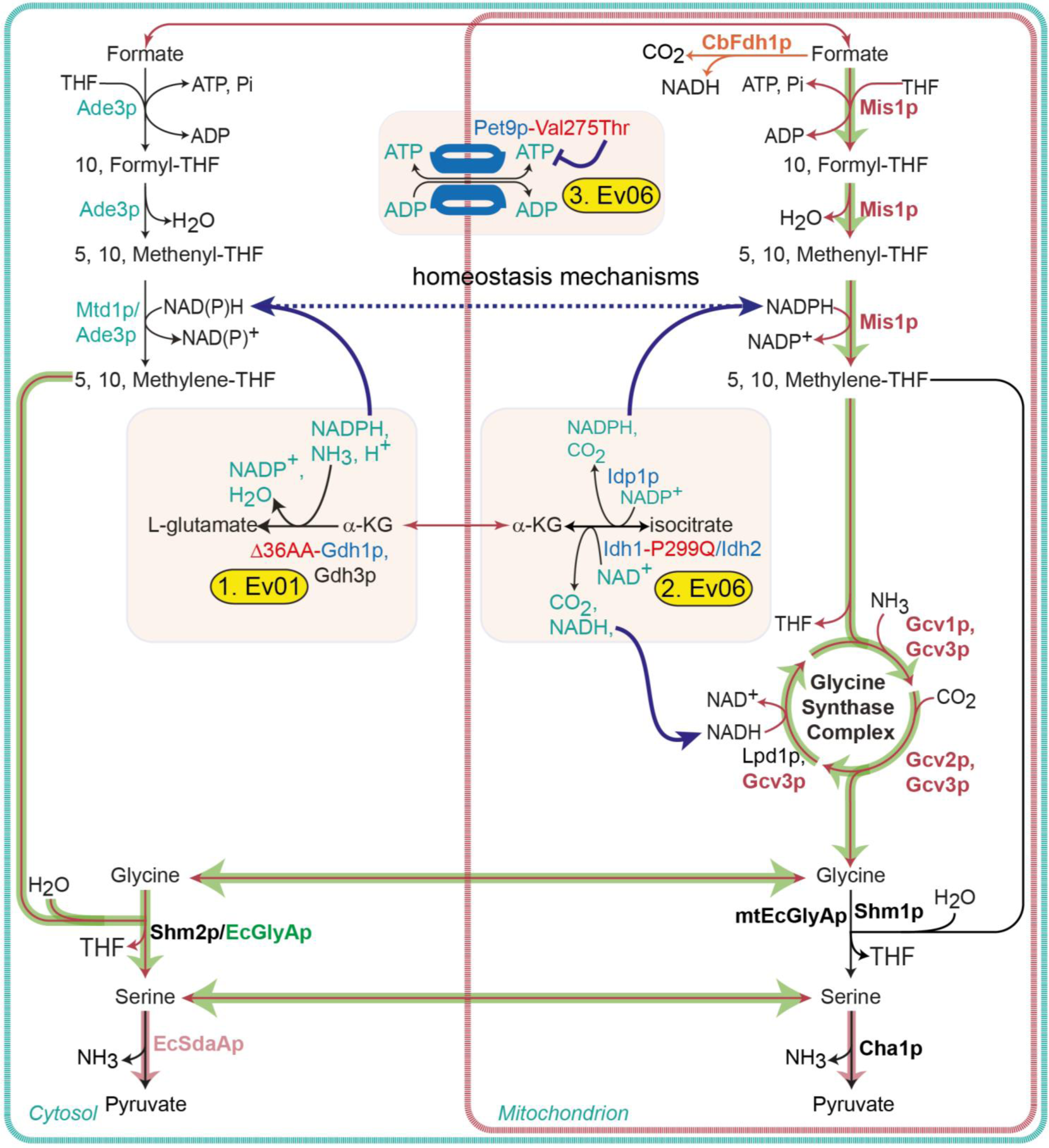
Potential effect of mutations in VBS10 on RGP activity. The glycine synthesis module is localized to the mitochondria. The glycine-to-serine conversion submodule composed of *Ec*GlyAp is localized to the cytoplasm and employs CH_2_-THF synthesized via cytoplasmic Ade3p/Mtd1p. The potential role of mutations promoting the RGP activity is displayed. Future engineering of serine deamination modules in the mitochondria (Cha1p) and the cytoplasm (*Ec*SdaAp) and expression of energy module (cbFdh1p) targeted to the mitochondria is indicated as well.

The *gdh1*::Δ108bp mutation, appearing early on during ALE, could directly enhance cytoplasmic formate conversion to CH_2_-THF and promote the glycine to serine conversion. This might also lead to the balancing of mitochondrial C1-THF and NADPH pools, thereby indirectly enabling operation of the glycine synthesis module (**Figure 8**). In addition, the *gdh1*::Δ108bp mutation will disrupt the efficient ammonia assimilation by this enzyme and may increase intracellular NH_4_ levels, which could help to push the glycine synthesis (by Gcv1p). This mutation may act as a temporary cue to enable RGP activity, accordingly, it can be reconstituted in the unevolved strain and facilitate pathway operation. Assuming *gdh1*::Δ108bp leads to high NADPH, recreating a similar physiological conditions by overexpression additional enzymes that leads to increased mitochondrial NADPH levels, might enable growth via the RGP.

Though the *gdh1::Δ108bp* mutation helps VBS10 switch to the RGP for glycine and serine synthesis, the growth rate without further evolution is very low. In the subsequent ALE steps, the average DT of 10 colonies from Ev03 was ∼74 h, sharply dropping to ∼19.75 h doubling time by Ev06. At this stage of ALE, the VBS10 strain acquired two new point mutations, i.e., Pet9p-I275T and Idh1p-P299Q. We speculated that these mutations play a significant role in the redistribution of energy and reducing power, thus enhancing the RGP-dependent growth. Pet9p is the major ATP-ADP transporter in the mitochondrial inner membrane. The mutation identified in the study is located towards the mitochondrial matrix^34^. This mutation might have a role in regulating ATP availability in the mitochondria and the cytoplasm, balancing the ATP supply for formate ligation to the mitochondrial or cytoplasmic THF. This improves the discrete C1-fluxes in both compartments that may facilitate glycine synthesis via GSC in mitochondria and glycine-to-serine conversion in the cytoplasm via *EcGlyA* (**Figure 8**).

At the same time, the Idh1p-P299Q mutation in the activation subunit of the mitochondrial, hetero octameric isocitrate dehydrogenase (Idh1p+Idh2p) is present in one of the AMP binding domains of the Idh1p subunit. Idh2p is a catalytic subunit that catalytically binds to isocitrate, which is activated by allosteric regulation of Idh1p via AMP and non-catalytic binding of citrate^44^. This AMP interacting domain composed of Asn298-Pro299-Thr300 domain mutated to Asn298-Gln299-Thr300. The effect of this mutation is unclear. The change of proline to glutamate might expose the AMP-binding residue to facilitate an AMP-activation-like confirmation, promoting the catalytic activity of the Idh complex. Assuming a positive effect on the catalytic activity, this mutation would have a direct role in promoting isocitrate to α-KG and NADH conversion, either promoting GCS activity or glutamate synthesis in the mitochondria via NAD-specific Glt1p (Glutamate synthase). However, if this mutation reduces AMP-dependent allosteric activation, leading to a complete loss or reduction of Idh activity, the NADP-specific Idp1p might take over this biochemical reaction synthesizing α-KG and NADPH, which would improve the mitochondria pool of NADPH, leading to increased CH_2_-THF availability and facilitating glycine synthesis via GSC^45^. We speculate that both mutations described above have a role in the growth improvements between Ev03 to Ev06. However, both require future investigation to analyze their precise effect on the RGP operation via reverse engineering and further testing in the *Δidp1* or *Δidh1* genotypes.

In summary, the results presented here form an essential step towards engineering full formatotrophic growth in *S. cerevisiae*. We demonstrated complete RGP activity in yeast. The initial goal of the present study – establishing glycine and serine biosynthesis from formate and CO_2_ – was exceeded by showing pyruvate (alanine) synthesis from all the stages of evolution via endogenous Cha1p activity. The ΔS strains, that we constructed in this study, could also be used to establish growth on methanol, by including a methanol to formate oxidation module, as previously shown in *E. coli* and *P. putida*^16,21,46^. Although the first attempts to apply ALE for increased flux towards pyruvate failed, the outlook is very promising. We envision that a combination of overexpression of a serine deaminase (in mitochondria and cytoplasm), integration of fast heterologous formate dehydrogenase candidates, combined with evolution under a glucose limiting regime with formate in a chemostat might enable the strain to become fully formatotrophic. As the mutations identified in this study indicate a high demand for energy and reducing power, it could be interesting to reverse engineer these mutations in strains to establish full formatotrophy, this may further speed up ALE. Another possible rational strategy would be to delete strong NADPH-consuming enzymes, such as the glyoxylate reductase Gor1p^47^, or the mitochondrial overexpression of the NAD^+^ dependent formate dehydrogenase (Fdh1p)^48^ along with the mitochondrial NADH kinase Pos5p^49^ in order to increase NADPH availability in the mitochondria^50,51^ that seems to play an important role in enabling the yeast-RGP.

## Authorship contribution

**Viswanada R Bysani (V.R.B.):** All experiments, analysis and interpretations were performed by V.R.B under the supervision of A.B.E. and F.M. Involved in supervision of students, visualization, investigation, data curation, interpretation, writing & editing.

**Ayesha Syed Mamoor Alam (A.S.M.A):** Involved in screening serine biosensor strains i.e., VBS08, VBS09, and VBS10 strain with *SHMT*ox and labeling data analysis under supervision V.R.B.

**Fabian Machens (F.M.): S**upervision, writing & editing.

**Arren Bar-Even (A.B.E.):** Project conceptualization, supervision (until his demise), Funding acquisition.

## Abbreviations/Keywords

RGP/rGlyP: Reductive Glycine Pathway
rAcCoAP: Reductive Acetyl-CoA pathway
GCS: Glycine Cleavage System (GCV system or GDC-Glycine Decarboxylase Complex)
GSC: Glycine Synthase Complex (Reversal of GCS)
CHO-THF: 10-formyltetrahydrofolate
CH-THF: 5,10-methenyltetrahydrofolate
CH_2_-THF: 5,10-methylenetetrahydrofolate
CH_3_-THF: 5-methyltetrahydrofolate
C1 units: CHO-THF, CH-THF, CH_2_-THF, and CH_3_-THF
3-PGA: 3-phosphoglyceric acid
SHMT: Serine hydroxymethyltransferase
ΔS: Serine biosensor
Ev: Evolution round
ALE: Adaptive Laboratory Evolution
NGS: Next Generation Sequencing
CDS: Coding sequence
ARS: Autonomously replicating sequence

## Acknowledgements

We thank Änne Michaelis for her help with the LC-MS analysis of amino acids, Dr. Vijay Jayaram for helping to understand *GDH1* and *IDH1* mutations and Dr. Nico Claassens for reading manuscript and giving critical comments.

## Materials and Methods

### Strains used in this study

*Saccharomyces cerevisiae* S288c (*MATα SUC2 gal2 mal2 mel flo1 flo8-1 hap1 ho bio1 bio6*) (a gift from Dr. Patrick Yizhi Cai’s lab, Manchester, UK) was used as the base strain in this study. All other yeast strains were derived from this parental strain are listed in the table (**Supplementary table 1**). For cloning and plasmid propagation, *E. coli* DH5α and NEB10ß cells (New England Biolabs GmbH, Frankfurt, Germany) were used.

### Media and growth conditions

YPD medium composed of 1 % (w/v) yeast extract, 2 % (w/v) peptone, and 2 % (w/v) glucose was used for the general culturing of yeast cells for DNA transformations and genome manipulations. 2x YPD (2 % yeast extract, 4 % peptone, 4 % glucose) was used for optimal growth prior to transformations. Synthetic Complete (SC) medium is composed of 1x (1.7 g/L) yeast nitrogen base (YNB) (Sigma-Aldrich, Steinheim, Germany), a customized Drop-Out mix of amino acids and nucleotides for optimal growth lacking the components used for selection e.g., Glycine and Serine in SC-Gly-Ser medium (), 5 g/L of ammonium sulfate ((NH_4_)_2_SO_4_) as nitrogen source unless differently indicated, and appropriate carbon sources. Synthetic Minimal (SM) medium comprises 1x YNB, 5 g/L (NH_4_)_2_SO_4_, and appropriate carbon sources. If not indicated otherwise, in both SC and SM media, 100 mM glucose was used as the primary carbon source, and 0.5 mM glycine, 0.5 mM serine, and 250 mM sodium formate were used as secondary carbon sources as indicated. All individual amino acids, nucleotides, and ammonium sulfate were procured from either Sigma-Aldrich or Carl Roth (Karlsruhe, Germany). All antibiotics used in yeast cultures or *E. coli* cultures were procured from Carl Roth. Yeast extract, peptone, glucose, sodium formate, ethanol, and isopropanol were procured from Carl Roth (Karlsruhe, Germany). Yeast strains were incubated at 30°C at 200 rpm and provided with an ambient or high CO_2_ atmosphere. Geneticin (G418) (200 mg/L), hygromycin (200 mg/L), and nourseothricin (100 mg/L) were used in yeast cultures for the selection of plasmids with KanR, HygR, and NatR cassettes. *E. coli* was always grown on LB medium (1% tryptone, 0.5% yeast extract, 1% NaCl) with appropriate antibiotic selection i.e., ampicillin (100 mg/L) or kanamycin (100 mg/L).

### PCR and DNA preparation

PCR reactions were performed with PrimeSTAR MAX DNA Polymerase (BD Clontech GmbH, Heidelberg, Germany) or Phusion High-Fidelity polymerase (Thermo Fisher Scientific GmbH, Dreieich, Germany) following the manufacturer’s recommendations. Colony PCR of *E. coli* cells was done with DreamTaq polymerase (Thermo Fisher Scientific GmbH, Dreieich, Germany). Phire Plant Direct PCR Master Mix (Thermo Fisher Scientific GmbH) was used for yeast colony PCR. All primers were synthesized by Eurofins Genomics GmbH (Ebersberg, Germany) or IDT (Leuven, Belgium) (For information on primers, refer to **Supplementary table 6** and Supplementary **Plasmid map 1-6**).

Lucigen Master pure (Lucigen Corp., WI, USA) and YeaStar Genomic DNA Kit (Zymo Research Freiburg, Germany) were used for genomic DNA isolation from yeast cells. Yeast plasmid miniprep II (Zymo Research) was used for plasmid preparation from yeast. GeneJET plasmid-miniprep-kit (Thermo Fisher Scientific GmbH) was used for plasmid preparations from *E. coli*. GeneJET PCR Purification Kit and GeneJET Gel-extraction kit (both Thermo Fisher Scientific) were used for DNA purifications.

### Plasmids

The pWS series of plasmids were gift from Tom Ellis (Addgene plasmid # 90516; http://n2t.net/addgene:90516; RRID:Addgene_90516) (**Supplementary table 2**). pWS082 is the guide-RNA entry vector used for assembling short gRNA oligos (20 bp). pWS173-KanR, pWS174-NatR, and pWS175-HygR carry the Cas9 expression cassette. These plasmids were used for sequential gene deletion of *AGX1*, *GLY1*, and *SER1* genes to construct serine biosensor strains. Refer to **Supplementary table 2** for information on all the plasmids used in this study.

The entire pCfB series of the plasmids are part of the EasyClone collection, were a gift from Irina Borodina (Addgene plasmid # 78231; http://n2t.net/addgene:78231; RRID:Addgene_78231) (**Supplementary table 2**). The pCfB2312 plasmid with KanR carries a Cas9 expression cassette under the *TEF1* promoter and *CYC1* terminator. The pCfB3041, pCfB3042, and pCfB3045 plasmids have sgRNA expression cassettes targeting the ChrX-3, ChrX-4, and ChrXI-3 loci, respectively, and a NatR selection cassette. The pCfB3034, pCfB3035, and pCfB2904 plasmids are genome integration vectors that carry ChrX-3, ChrX-4, and ChrXI-3 homology arms^52^. These plasmid backbones were amplified and used to construct genome integrative donor DNAs of the pathway modules.

The pVB05 plasmid is a genome integration vector with homology regions for the ChrX-4 locus made from the pCfB3035 backbone. The *SHM1* overexpression module, under *PGK1* promoter and terminator, was amplified using pLH52 as a template and assembled between ChrX-4 homology regions in the pCfB3035 plasmid backbone to construct the pVB05 plasmid (**Plasmid map 1**). Similarly, the *SHM2* overexpression module, under *GMP1* promoter and terminator, was amplified from the pLH53 template and cloned in pCfB2904 between the ChrXI-3 homology regions to create pVB06 plasmid (**Plasmid map 2**). To construct the pVB07 plasmid (**Plasmid map 3**), the *EcGlyA* expression cassette under the control of the *RPL3* promoter and *CWP2* terminators was amplified from the pVB01 plasmid (**Plasmid map 4**) and assembled in the pCfB3035 plasmid between the homology regions for genome integration into ChrX-4 locus. These plasmids were constructed using the NEBuilder HiFi DNA assembly master mix as per the supplier’s recommendation (New England Biolabs GmbH, Frankfurt, Germany). The pLH52 and pLH53 plasmids were a gift from Dr. Lena Hochrein (University of Potsdam, Germany).

The pFM340 plasmid (**Plasmid map 5**) carries expression cassettes *MIS1*, *GCV1*, *GCV2*, and *GCV3* of the glycine synthesis module of the RGP. The pFM340 plasmid was created by replacing the *URA3* selection marker on the original pJGC3^25^ (**Plasmid map 6**) plasmid with a HygR cassette.

### Yeast transformation

DNA transformations into yeast were performed using the lithium acetate-based heat shock method^53,54^. Recovered transformants were selected based on the selection marker available, and colonies were screened for mutations by PCR amplification and Sanger sequencing of the amplified locus. Important primer used for screening and sequencing are listed in the Supplementary information (**Supplementary table 6**).

### Strain engineering

The serine biosensor (ΔS) strain, VBS01 (S288c_ΔS), was generated in the *S. cerevisiae* S288c background by making knockouts of *AGX1*, *GLY1*, and *SER1* genes to block the serine and glycine routes (**Table 1**). All knockouts were verified by PCR and Sanger sequencing (at LGC Genomics, Berlin, Germany). To engineer further modules of the RGP in the serine biosensor strain, pCfB2312-Cas9 was transformed into VBS01 to create VBS03. The VBS04 strain was constructed by genome integration of the *SHM1ox* module into the ChrX-4 locus of the VBS03 strain. The *SHM1* module, with ChrX-4 homology arms, was amplified from pVB05 and transformed into VBS03 along with the pCfB3042-sgChrX-4 plasmid. Transformants were selected based on G418, and nourseothricin resistances were screened for the successful integration events by colony PCR and confirmed by Sanger sequencing.

VBS05 strain was constructed by integrating the *SHM2* module into the ChrXI-3 locus of VBS03. The *SHM2* overexpression module, with ChrXI-3 homology regions, was amplified from pVB06 and transformed into VBS03 along with the pCfB3045-sgChrXI-3 plasmid. The *EcGlyA* module, with ChrX-4 homology arms, was amplified from pVB07 and transformed into VBS03 along with the pCfB3042-sgChrX-4 plasmid to create VBS06 strain. These three ΔS strains, VBS04, VBS05, and VBS06, respectively, overexpress *SHM1ox, SHM2ox,* and *EcGlyAox* modules, involved in the glycine to serine conversion in the RGP. To further construct ΔS strains expressing the complete serine synthesis module, the VBS08, VBS09, and VBS10 strains were made by transforming the pFM340 plasmid (carries glycine synthesis module) into VBS04, VBS05, and VBS06 strains, respectively. Additionally, VBS18 and VBS19 strains were created by making a knockout of 108 bp from the ORF of the *GDH1* gene on genomes of VBS10 and S288c_WT strain, respectively.

### CRISPR-Cas9 methodology for gene deletions

For gene knockout using CRISPR-Cas9 methodology, pWS082, pWS173, pWS174, and pWS175 plasmids were used. This CRISPR-Cas9 methodology is optimized by combining principles and tools described in^55–57^ publications and in https://benchling.com/pub/ellis-crispr-tools. *SER1*, *AGX1*, and *GLY1* genes were deleted by making marker-free complete CDS knockouts.

The sgRNA oligos (**Supplementary table 6**) were designed using Yeastriction online tools^58^ (http://yeastriction.tnw.tudelft.nl/#!/) and Benchling CRISPR tool for the targeted genes (http://www.benchling.com), e.g., *SER1*, *AGX1*, *GLY1*, and *GDH1*. The GACTTT sequence is added to the upper oligo’s 5’ end of the upper oligonucleotides. CAAA is added to the 5’ end and AA to the 3’ end of the lower primer to complete the HDV ribosomal sequence for stability sgRNA^57,59^. Then both primers were annealed in a thermocycler by a stepwise decrease of temperature from 95°C to 25°C. The annealed double standard guide oligonucleotides with GACT and CAAA overhangs are assembled into the pWS082 vector^59^.

A 100 ng of pWS082, 2 µL of annealed oligos with overhangs, 1 µL of T4 DNA ligase, and Esp3I restriction enzyme were mixed in 1x T4 ligase buffer. The combined restriction digestion/ligation was performed in a thermocycler: 30 x (5min at 30°C, 5 min at 16°C)^55^. Subsequently, this Golden-Gate reaction was transformed into competent *E. coli* DH5α and selected for ampicillin-resistant transformants. Non-GFP colonies were picked as positive clones considering the GFP expression cassette in the pWS082 is replaced by sgRNA oligonucleotide insertion. The plasmid was prepared from these non-GFP colonies, and oligonucleotide insertion was confirmed by sequencing. The verified pWS082 plasmids with gDNA inserts were used as templates to amplify the whole sgRNA expression cassettes for transformation.

The Cas9 expression plasmids, i.e., pWS173 with KanR, pWS174 with NatR, and pWS175 with HygR, carry a 2µ Ori, a nuclear targeted Cas9 expression cassette, and a super-folder *GFP* expression cassette cloned between Esp3I restriction sites. To prepare Cas9 backbones with different selection markers, these plasmids were digested with Esp3I, and the backbone DNA fragment containing the Cas9 cassette and the respective selection marker were gel purified. This fragment carries homology sequences, at both ends, to the amplified sgRNA expression cassettes from pWS082 vectors.

To swap the targeted gene or partial DNA sequence from the genome, repair DNA fragments were designed using Yeastriction online tool, SnapGene 6.02 (from Insightful Science; available at snapgene.com), or Geneious 11.1.5 software (https://www.geneious.com). A 60 bp of the upstream region and a 60 bp of the downstream region of the targeted deletion were combined separately from both DNA strands and generated a pair of 120 bp long primers complementing each other. The primers were annealed and used as double-stranded donor DNA for genome repair. If necessary, point mutations were introduced to destroy PAM sequences and prevent the potential Cas9 binding to the already edited locus or the free double-stranded donor DNA fragments.

The linear sgRNA fragment, linear Cas9 backbone, and linear donor/repair DNA fragments were transformed in the intended yeast strain and selected according to the resistant cassettes. Due to the mutual homology sequences, the sgRNA fragment and Cas9 fragment form a single circular plasmid by activating the native recombination machinery of yeast^60^. These transformants can be selected based on the antibiotic resistance cassette as the circular plasmid express resistance marker, Cas9 protein, and sgRNA. The tRNA fragment of sgRNA undergoes self-cleavage upon expression, leaving a mature, stable, persistent sgRNA on which Cas9 is assemble. The sgRNA and Cas9 complex introduces a double standard break in the genome, which are repaired by already activated recombination machinery according to the homology regions on the transformed donor DNA.

### Adaptive Laboratory Evolution

Adaptive Laboratory Evolution (ALE) was conducted with VBS08, VBS09, and VBS10 strains in the SC-Gly-Ser medium. SC medium used for ALE is composed of 1x YNB, 100 mM (NH_4_)_2_SO_4_, 100 mM glucose, and 100 or 250 mM formate. SC medium contains a Drop-Out mix of amino acids, including standard concentrations of uracil or adenine, in defined quantities (Sigma-Aldrich, 2014-Y1751 & Y1856). A custom mix of Drop-Out without serine and glycine was prepared and used in the SC-Gly-Ser medium to establish glycine and serine synthesis from formate using growth as a readout. ALE was continued in SM medium composed of 1x YNB, 100 mM (NH_4_)_2_SO_4_, 100 mM glucose, and 100 or 250 mM formate. The maintenance of the glycine synthesis module of the RGP encoded on pFM340 was achieved with hygromycin selection (200 mg/L).

### Determination of growth improvements in the ALE

The growth improvements of VBS10 strains, operating the RGP for the formate-dependent synthesis of serine and glycine were assessed through high throughput growth experiments. The growth rates of VBS10 strains from Ev03, Ev06, Ev12, Ev16, and Ev21 were evaluated by screening multiple colonies from each selected evolution round using multi-well readers. The glycerol stocks of the selected evolution rounds were streaked on YPD hygromycin plates and incubated at 30°C until prominent colonies appeared. 10 colonies from each selected evolution round were picked and inoculated in SM medium with 50 mM glucose and 250 mM formate for primary culture preparation. These cultures were incubated under a 10% CO_2_ atmosphere at 30°C with 200 rpm of orbital shaking until growth appeared. The precultures were washed in sterile distilled water and used as seed culture to determine the growth rate in SM medium with 50 mM glucose with or without 100 mM formate. All the 10 cultures were inoculated to OD_600_ = 0.02 in 1.5-2 ml Eppendorf tubes in the selective medium and mixed. The homogenous cultures were distributed in 48 well plates in technical duplicates. To prevent evaporation during the experiment, each well was filled with a culture volume of 350 µl and overlayered with 250 µl of mineral oil. The plates were incubated in the multi-well plate reader instruments (Tecan – Spark/infinite, Crailsheim, Germany or BioTek Epoch 2 - BioTek, Bad Friedrichshall, Germany) at 30 °C, providing a 10% CO_2_ atmosphere. Alternating orbital and linear shaking (∼300 rpm, 60 sec per shaking mode) was programmed, and growth was measured at ∼15 mins intervals by reading optical density at 600 nm wavelength in each cycle. The output data of the optical density values were used to plot growth curves using a custom MATLAB script (available on request) (MATLABR2019a). In the MATLAB script, the OD values of the 48 well plates were extrapolated to cuvette OD values using a predetermined conversion factor. The MATLAB script also calculates the growth rate and doubling time. The growth rates of multiple colonies from selected evolution rounds were plotted and analyzed by one-way ANOVA followed by Tukey multiple comparisons tests using GraphPad Prism version 9 for Windows (GraphPad Software, San Diego, California USA, www.graphpad.com).

### Stable ^13^C isotope tracing of formate and CO_2_ utilization

To confirm the metabolic pathway responsible for carbon assimilation from formate to glycine, serine, and pyruvate in the evolved VBS10 strains, stable ^13^C isotope labelling experiments were conducted with the evolved VBS10 strains from different evolution rounds. The VBS10 strains from Ev02, Ev06, Ev10, Ev16, Ev21, Ev25, and Ev29 were cultured in the SM medium supplied with ^13^C or ^12^C formate, with 40 mM glucose as the primary carbon source. The cultures were incubated in a 10% ^13^CO_2_ or ^12^CO_2_ atmosphere. A vacuum desiccator (Lab Companion, MA, USA) was used to grow cultures in ^13^CO_2_, where the original gas was drawn-out by a vacuum pump, followed by refilling with 10% ^13^CO_2_ and 90% air. Cells equivalent to 2 OD_600_ from grown cultures were collected from the early stationary phase and centrifuged at 20,000 x g for 5 mins to pool the cell biomass. The washed biomass was dissolved in 1 ml 6 N HCL and hydrolyzed overnight at 95 °C. Then samples were dried at the same temperature with or without flushing air^61^. After drying the samples at 95 °C, the samples were resuspended in ultra-pure water, filtered, and the masses of amino acids were analyzed with UPLC-ESI-MS as previously described^62^. Hydrolyzed proteinogenic amino acids were separated using the Waters Acquity UPLC system (Waters, Milford, MA, USA), equipped with an HSS T3 C18 reversed-phase column (100 x 2.1 mm2, 1.8 mm; Waters, Eschborn, Germany). In the mobile phase, 0.1% formic acid in H_2_O (A) and 0.1% formic acid in acetonitrile (B) were used. The flow rate was 0.4 mL/min, and the gradient was: 0 to 1 min – 99% A; 1 to 5 min – linear gradient from 99% A to 82%; 5 to 6 min – linear gradient from 82% A to 1% A; 6 to 8 min – kept at 1% A; 8 to 8.5 min – linear gradient to 99% A; 8.5 to 11 min – re-equilibrate. Mass spectra were acquired using an Exactive mass spectrometer (ThermoScientific, Dreieich, Germany) in positive ionization mode, with a scanning range of 50.0 to 300.0 m/z. Spectra were recorded during the first 5 min of the LC gradients. Under these conditions, the retention times for amino acids were determined by analyzing amino-acid standards (Sigma-Aldrich) under the same conditions. Data analysis was performed using Xcalibur (ThermoScientific, Dreieich, Germany).

### Genome sequencing analysis

Genomic DNA samples were run through an additional RNase treatment to clean the contaminant RNA before sending for whole-genome sequencing. For this, samples resuspended in TE buffer and RNase were added and incubated at 37°C for one hour ^63^. Then samples were precipitated and resuspended in TE buffer to send for NGS. Library construction of genomic DNA samples and sequencing was performed by Novogene (Cambridge, United Kingdom) using the paired-end Illumina sequencing platform. The sequencing data was analysed using the breseq pipeline^64^ and modified for multi chromosomal haploid organisms. The S288c genome sequences downloaded from the NCBI database were used as the reference genome to compare the genome changes acquired in the ALE. A non-modified wild-type strain was sequenced along with the evolved and unevolved strains as a control. NGS data of WT, unevolved VBS10, and evolved VBS10 strains were aligned to the S288c reference sequence and compared mutations. Genome sequencing was done for 4 biological replicates from each selected evolution round of VBS10. All mutations, that occurred in 100% of reads in all four replicates (four biological replicates) from each evolution round, were considered as mutations that might support formate-dependent growth improvement directly or indirectly. The list of mutations identified was segregated based on their location, i.e., either on the coding sequences or in the intergenic regions of genomes.

## Data availability

Additional information on the experimental setup as well as detailed results are available from the corresponding author upon request. Any strains and plasmids generated during this study are available upon completing a Materials Transfer Agreement.

## Code availability

MATLAB and breseq codes used for the analysis of the experiments are available from the corresponding author upon request.

## Supplementary information

**Supplementary figure 1: Phenotype analysis of VBS01 strain: A.** Growth of VBS01 strain in SM medium supplemented with glycine/serine in a 10% CO_2_ atmosphere along with S288c_WT control without serine or glycine supplementation. **B.** Growth curves of VBS01 strain in SM+formate medium supplemented with glycine or serine in a 10% CO_2_ atmosphere along with WT control without serine or glycine supplementation. **DT:** Doubling Time in hours; **NG:** No Growth;

**Supplementary figure 2: Growth curves of VBS10 strain from Ev01 and Ev04 in SM medium: A.** Growth of the VBS10_Ev01 in SM medium with or without formate, supplied with 0.5 mM glycine. All cultures with different media conditions were incubated in a 10% CO_2_ atmosphere. **B.** Growth of the VBS10_Ev04 strain in SM medium supplied with formate and incubated in either 10% CO_2_ or ambient CO_2_ condition. Growth is observed only when formate is supplied as the secondary carbon source and incubated under a 10% CO_2_ atmosphere. **SM:** 1xYNB, 100 mM (NH_4_)_2_SO_4_, and 100 mM glucose unless stated otherwise; **YNB:** Yeast Nitrogen Base without (NH_4_)_2_SO_4_.

**Supplementary figure 3:** Metabolic schemes of the distribution of the ^13^C into glycine, serine, pyruvate, and alanine when ^13^C carbon was supplied via formate and CO_2_ combinations.

**Supplementary figure 4:** ^13^C-formate labeling data shows that nearly all one-carbon units are generated from ^13^C-formate in the evolved VBS10 strain, as formate-derived C_1_-units are assimilated to histidine and methionine. WT assimilates relatively less ^13^C into histidine and methionine, compared to WT. However, the date shows the efficient reversible activity of Ade3p/Mis1p.

**Supplementary figure 5: Evolution of VBS10_Ev21 strain in glucose limiting conditions:** The evolved VBS10_Ev21 strain further evolved in decreasing glucose conditions. Growth curves of the VBS10 in 40 mM, 20 mM, 10 mM, 5 mM, and 2 mM glucose, respectively, in the Ev22, Ev25, Ev27, Ev28, and Ev29 in SM medium supplemented with 250 mM and 100 mM formate.

**Supplementary figure 6: Formate, CO_2_, and (NH_4_)_2_SO_4_ dependency of the VBS10_Ev21 strain: A.** Growth of VBS10_Ev21 strain in various concentrations of formate as the secondary carbon source ranging from 2 mM to 1 M while incubated under 10% CO_2_. **B.** Growth of the VBS_Ev21 strains when incubated in different CO_2_ atmospheres, i.e., 1%, 2.5%, 5%, and 10% CO_2_ atmospheres with 100 mM formate across the experiments. **C.** Growth of the VBS_Ev21 and D. S288c_WT strains in various concentrations of (NH_4_)_2_SO_4_ while formate and CO_2_ are maintained constant.

**Supplementary figure 7: Idh1p::P299Q in the AMP binding domain:** Amino acids from 298-300 are the AMP binding domain of the IDH complex. The identified point mutation changes the AMP binding domain from Asn298:Pro299:Thr300 to Asn298:Gln299:Thr300. The proline (hydrophobic) change to glutamine (hydrophilic) might apart the interaction between Asn:Thr and change to the confirmation to a similar state as when the Idh complex is bound to AMP.

**Supplementary figure 8: Reconfirmation of evolution with reverse engineering of the *GDH1* knockout:** Growth of the VBS10_Δgdh1::108bp isolates in comparison with VBS10in SC-Ser-Gly medium with 100 mM glucose and 250 mM formate. All four (C1-C4) clones of VBS10_Δgdh1::108bp grew in 25 days, whereas both VBS10 clones did not grow until 40 days.**SC-Gly-Ser:** 1x YNB, DO-Gly-ser, 100 mM (NH_4_)_2_SO_4_, 100 mM glucose, 250 mM formate. **DO-Gly-Ser:** Dropout mix of all the amino acids and required nucleotides except serine and glycine.

**Supplementary figure 9:** Differences of ^13^C_1_-histidine, ^13^C_1_-methionine, and ^13^C_1_-serine between VBS19 (S288c *gdh1::Δ108bp*) and its WT controls. Δgdh1 = *gdh1::Δ108bp*; Y-scale split. T-test was performed, and the p-value and the significance were indicated.

### Supplementary plasmid maps with primers

**Plasmid map 1:** Map of pVB05 plasmid map with *SHM1* expression module under the *PGK1* promoter and terminator cloned between ChrX-4 integrable homology arms.

**Plasmid map 2:** Map of pVB06 plasmid with *SHM2* expression module under the *GPM1* promoter and terminator cloned between ChrXI-3 integrable homology arms.

**Plasmid map 3:** Map of pVB07 plasmid with *EcGlyA* expression module under the *RPL3* promoter and *CWP2* terminator cloned between ChrX-4 integrable homology arms.

**Plasmid map 4:** Map of pVB01 plasmid with *EcGlyA* expression module under the *RPL3* promoter and *CWP2* terminator.

**Plasmid map 5:** Plasmid map of pFM340 with *MIS1*, *GCV1*, *GCV2*, and *GCV3* genes under the promoter and terminators of *PYK1*, *TEF1*, *TEF2*, and *RPL3,* respectively. pFM340 carries a Hygromycin resistance cassette for selection in the yeast system. It is a centromere plasmid with CEN/ARS Ori.

**Plasmid map 6:** Plasmid map of pJGC3 with *MIS1*, *GCV1*, *GCV2*, and *GCV3* genes under the promoter and terminators of *PYK1*, *TEF1*, *TEF2*, and *RPL3,* respectively. pFM340 carries a *URA3* selection marker to maintain in the yeast system. It is a centromere plasmid with CEN/ARS Ori.

## Supplementary tables information

**Supplementary table 1: List of yeast strains and expression modules of the RGP used in this study. Supplementary table 2: List of plasmids used in this study:** Chr**-chromosome;**

**Supplementary table 3: List of strains and media conditions used for testing glycine and serine synthesis using VBS08, VBS09, and VBS10 in SC-Gly-Ser and SM media conditions: SC:** Synthetic Complete or semi-rich medium, **SM:** Synthetic minimal medium. **DO-Gly-Ser** is a mix of all the amino acids nucleotides in recommended quantities for optimal growth of yeast except serine and glycine, (**DO:** dropout);

**Supplementary table 4: List mutations identified from the NGS data VBS10 strains during ALE. Supplementary table 5: List of components used in DO-Ser-Gly mix.**

**Supplementary table 6: List of the primers used for screening and sequencing.**

