## Supplemental information for "Engineering and Evolution of the Complete Reductive Glycine Pathway in *Saccharomyces cerevisiae* for Formate and CO_2_ Assimilation"

### Supplementary information

### Supplementary figures

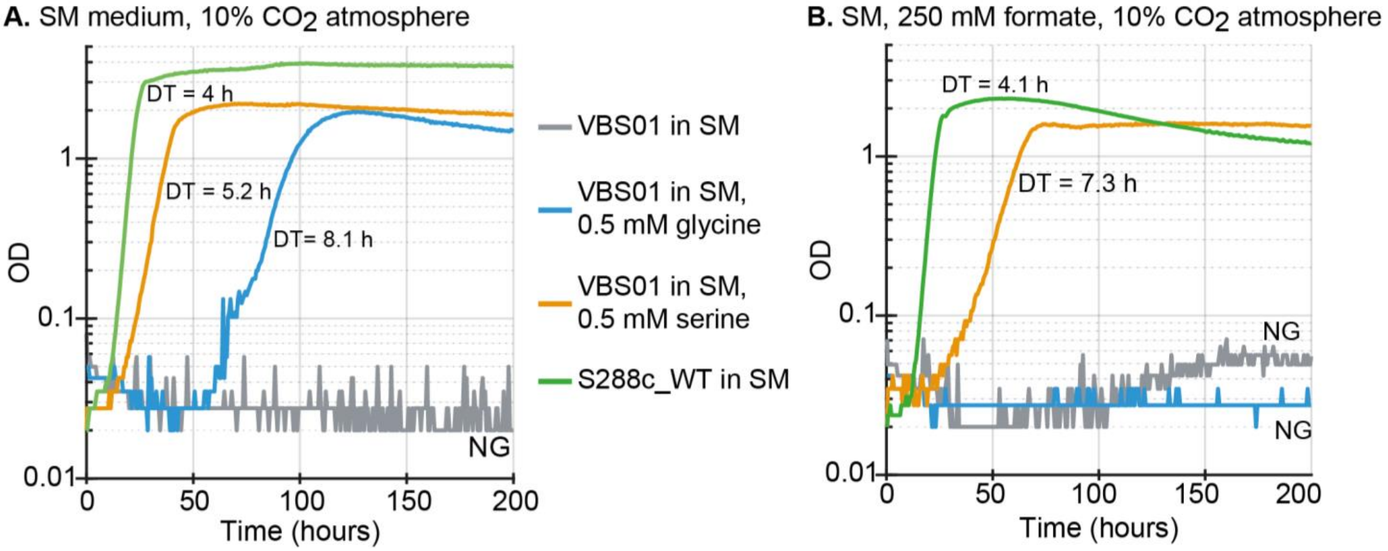


**Supplementary figure 1: Phenotype analysis of VBS01 strain: A.** Growth of VBS01 strain in SM medium supplemented with glycine/serine in a 10% $CO_2$ atmosphere along with S288c_WT control without serine or glycine supplementation. **B.** Growth curves of VBS01 strain in SM+formate medium supplemented with glycine or serine in a 10% $CO_2$ atmosphere along with WT control without serine or glycine supplementation. **DT:** Doubling Time in hours; **NG:** No Growth;

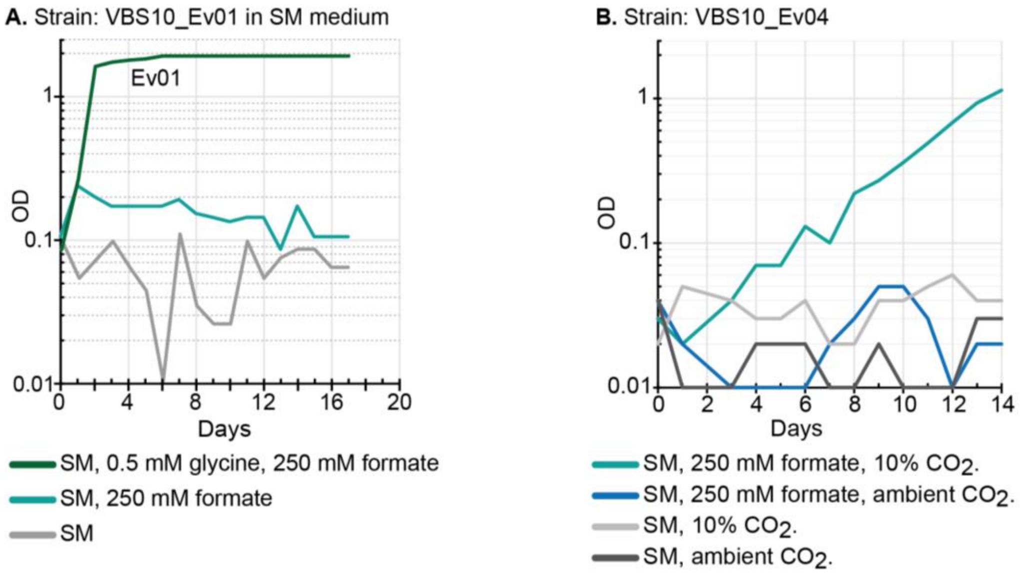


**Supplementary figure 2: Growth curves of VBS10 strain from Ev01 and Ev04 in SM medium: A.** Growth of the VBS10_Ev01 in SM medium with or without formate, supplied with 0.5 mM glycine. All cultures with different media conditions were incubated in a 10% $CO_2$ atmosphere. **B.** Growth of the VBS10_Ev04 strain in SM medium supplied with formate and incubated in either 10% $CO_2$ or ambient $CO_2$ condition. Growth is observed only when formate is supplied as the secondary carbon source and incubated under a 10% $CO_2$ atmosphere. **SM:** 1xYNB, 100 mM $(NH_4)_2SO_4$, and 100 mM glucose unless stated otherwise; **YNB:** Yeast Nitrogen Base without $(NH_4)_2SO_4$.

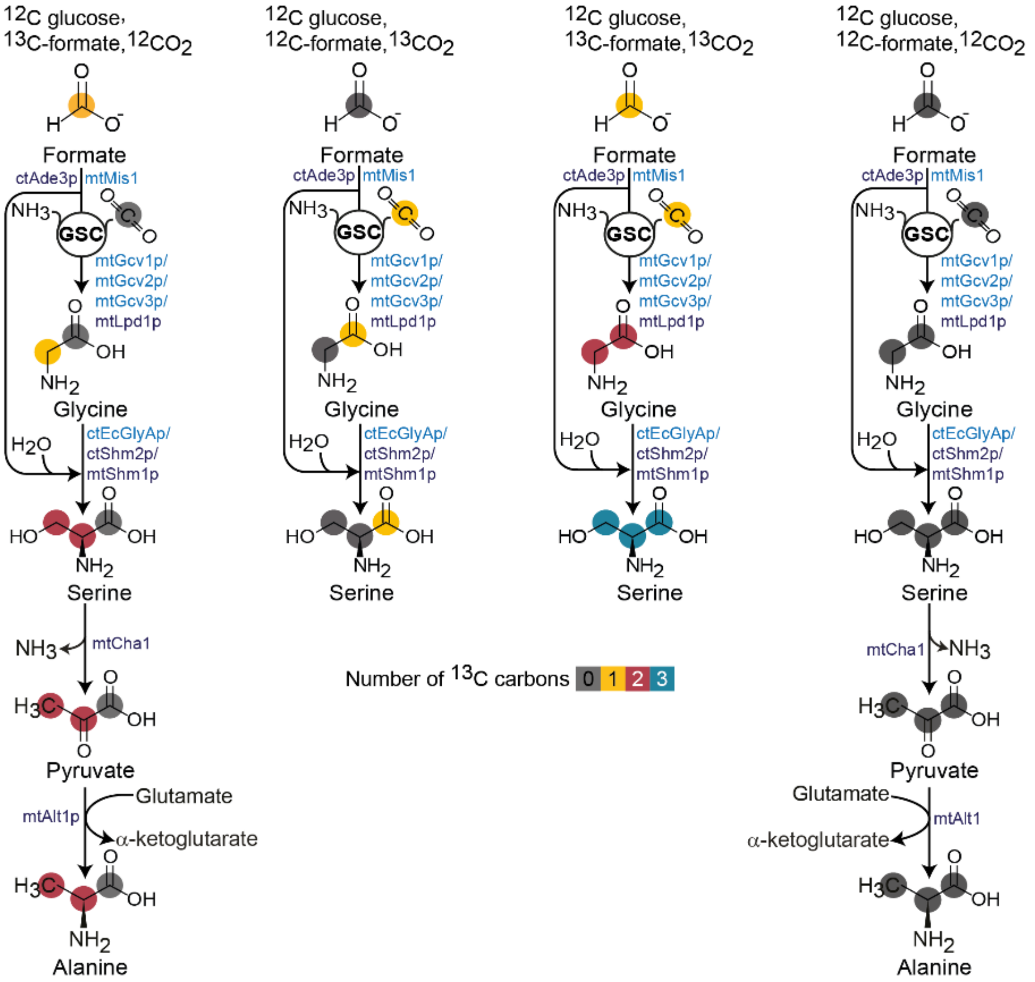


**Supplementary figure 3:** Metabolic schemes of the distribution of the $^{13}C$ into glycine, serine, pyruvate, and alanine when $^{13}C$ carbon was supplied via formate and $CO_2$ combinations.

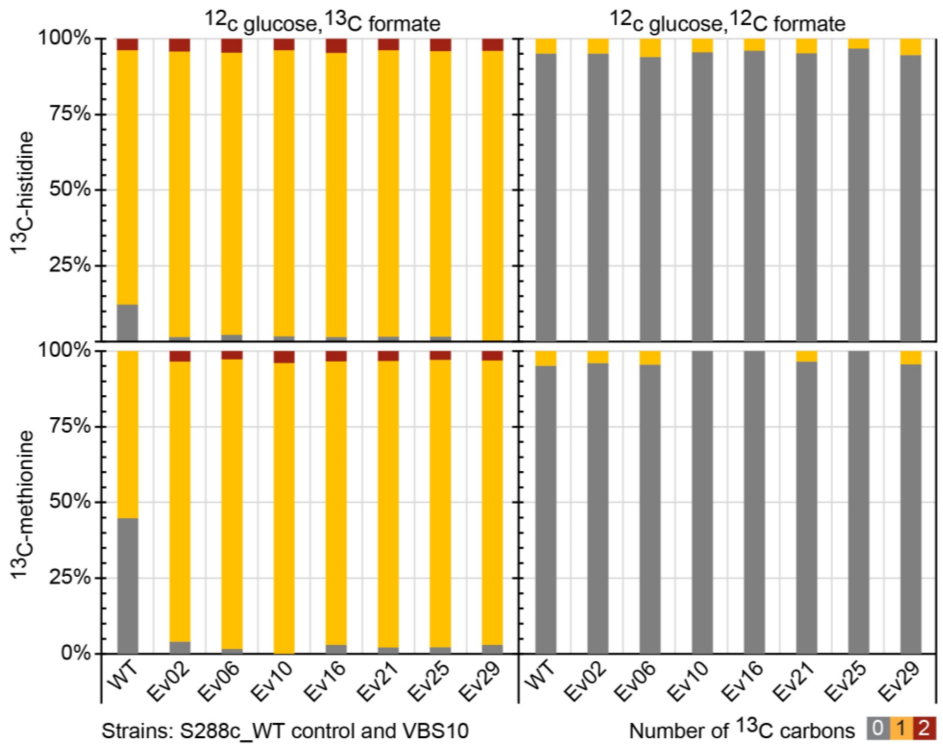


**Supplementary figure 4:** $^{13}C$-formate labeling data shows that nearly all one-carbon units are generated from $^{13}C$-formate in the evolved VBS10 strain, as formate-derived $C_1$-units are assimilated to histidine and methionine. WT assimilates relatively less $^{13}C$ into histidine and methionine compared to WT. However, the date shows the efficient reversible activity of Ade3p/Mis1p.

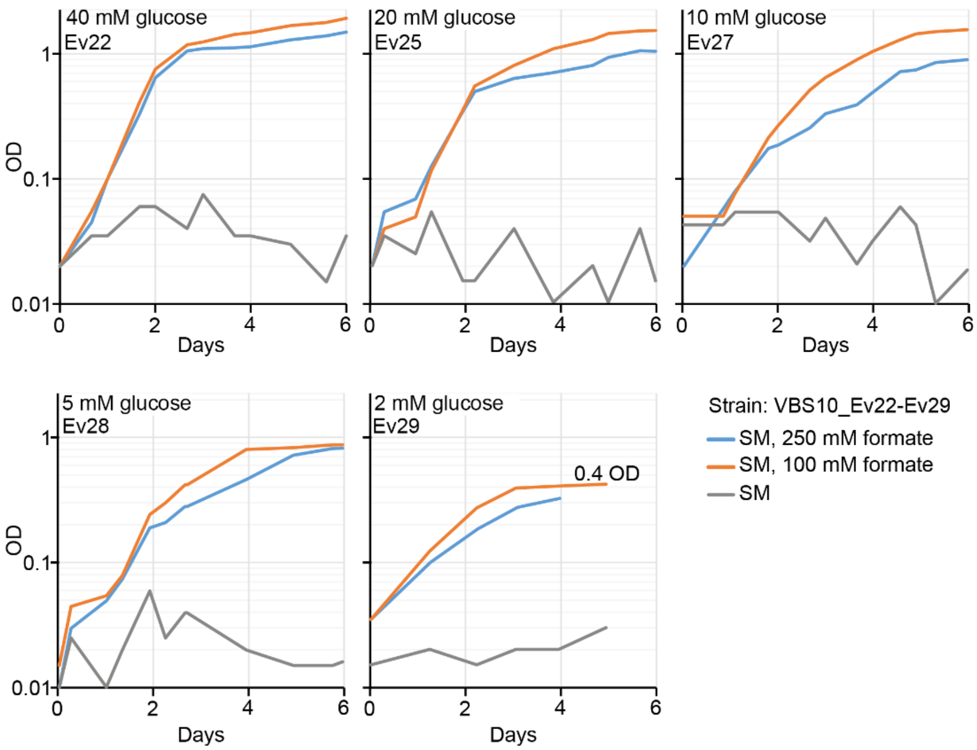


**Supplementary figure 5: Evolution of VBS10_Ev21 strain in glucose limiting conditions:** The evolved VBS10_Ev21 strain further evolved in decreasing glucose conditions. Growth curves of the VBS10 in 40 mM, 20 mM, 10 mM, 5 mM, and 2 mM glucose, respectively, in the Ev22, Ev25, Ev27, Ev28, and Ev29 in SM medium supplemented with 250 mM and 100 mM formate.

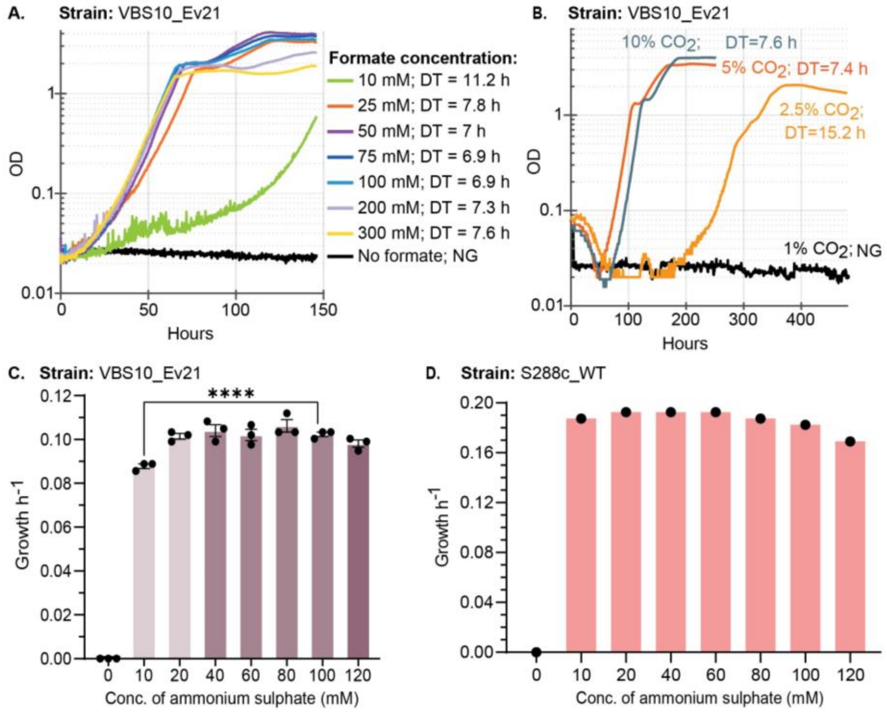


**Supplementary figure 6: Formate, $CO_2$, and $(NH_4)_2SO_4$ dependency of the VBS10_Ev21 strain: A.** Growth of VBS10_Ev21 strain in various concentrations of formate as the secondary carbon source ranging from 2 mM to 1 M while incubated under 10% $CO_2$. **B.** Growth of the VBS_Ev21 strains when incubated in different $CO_2$ atmospheres, i.e., 1%, 2.5%, 5%, and 10% $CO_2$ atmospheres with 100 mM formate across the experiments. **C.** Growth of the VBS_Ev21 and D. S288c_WT strains in various concentrations of $(NH_4)_2SO_4$ while formate and $CO_2$ are maintained constant.

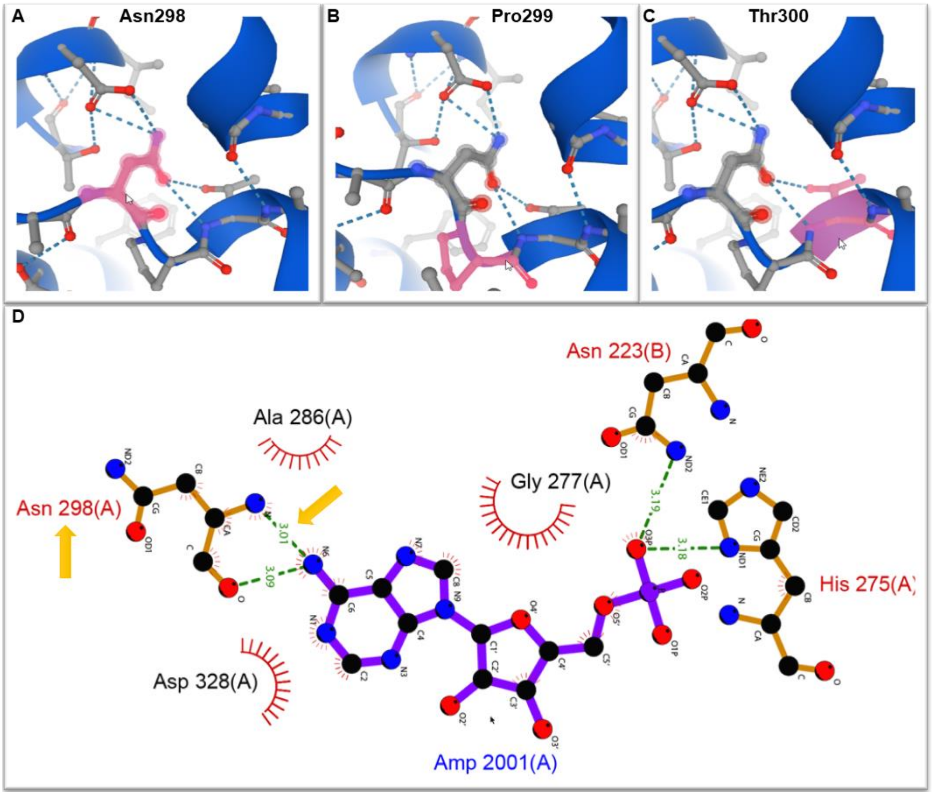


**Supplementary figure 7: Idh1p::P299Q in the AMP binding domain:** Amino acids from 298-300 are the AMP binding domain of the IDH complex. The identified point mutation changes the AMP binding domain from Asn298:Pro299:Thr300 to Asn298:Gln299:Thr300. The proline (hydrophobic) change to glutamine (hydrophilic) might apart the interaction between Asn:Thr and change to the confirmation to a similar state as when the Idh complex is bound to AMP.

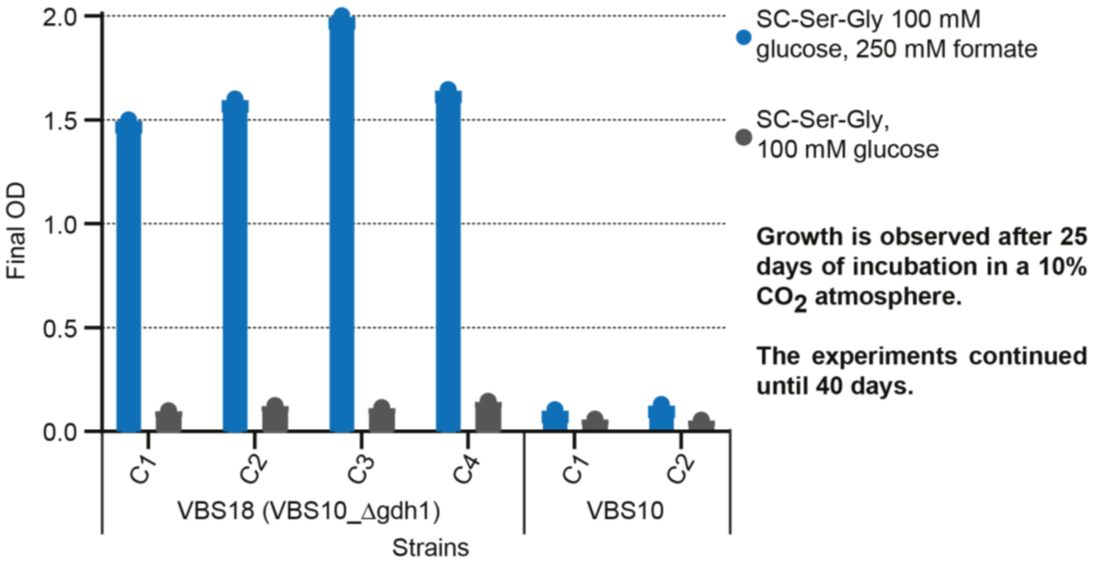


**Supplementary figure 8: Reconfirmation of evolution with reverse engineering of the *GDH1* knockout:** Growth of the VBS10_Δgdh1::108bp isolates in comparison with VBS10in SC-Ser-Gly medium with 100 mM glucose and 250 mM formate. All four (C1-C4) clones of VBS10_Δgdh1::108bp grew in 25 days, whereas both VBS10 clones did not grow until 40 days.**SC-Gly-Ser:** 1x YNB, DO-Gly-ser, 100 mM $(NH_4)_2SO_4$, 100 mM glucose, 250 mM formate. **DO-Gly-Ser:** Dropout mix of all the amino acids and required nucleotides except serine and glycine.

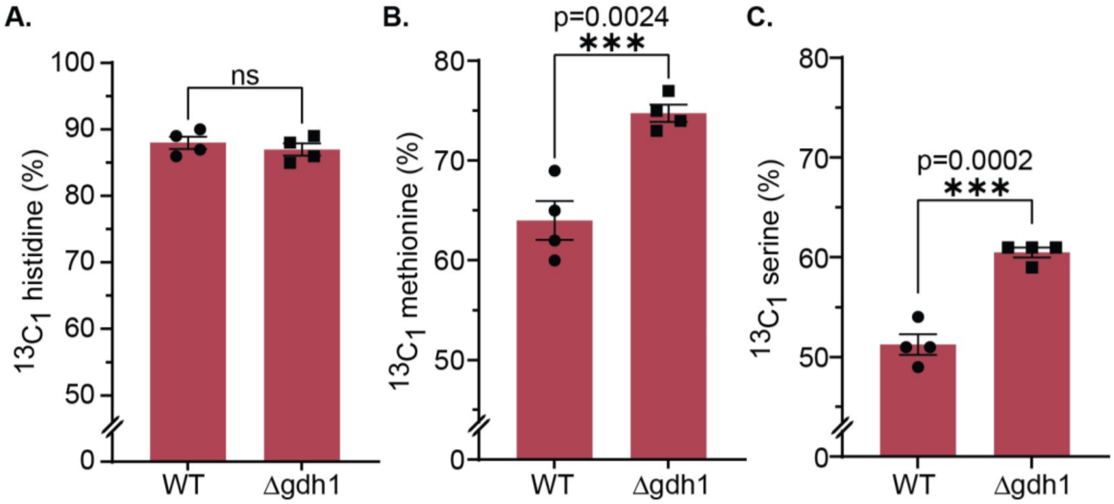


**Supplementary figure 9:** Differences of $^{13}C_1$-histidine, $^{13}C_1$-methionine, and $^{13}C_1$-serine between VBS19 (S288c *gdh1::Δ108bp*) and its WT controls. Δgdh1 = *gdh1::Δ108bp*; Y-scale split. T-test was performed, and the p-value and the significance were indicated.

### Supplementary plasmid maps with primers

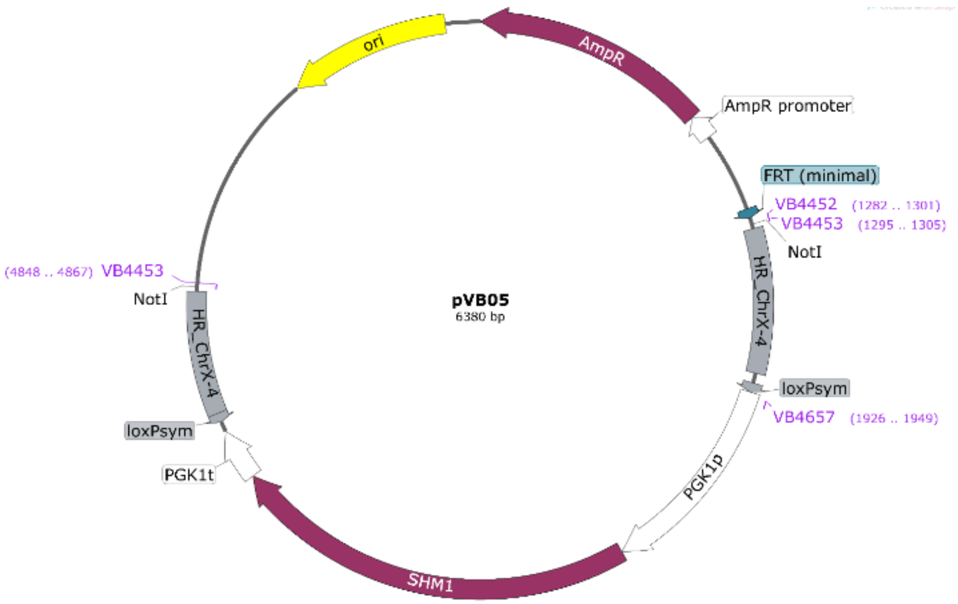


**Plasmid map 1:** Map of pVB05 plasmid map with *SHM1* expression module under the *PGK1* promoter and terminator cloned between ChrX-4 integrable homology arms.

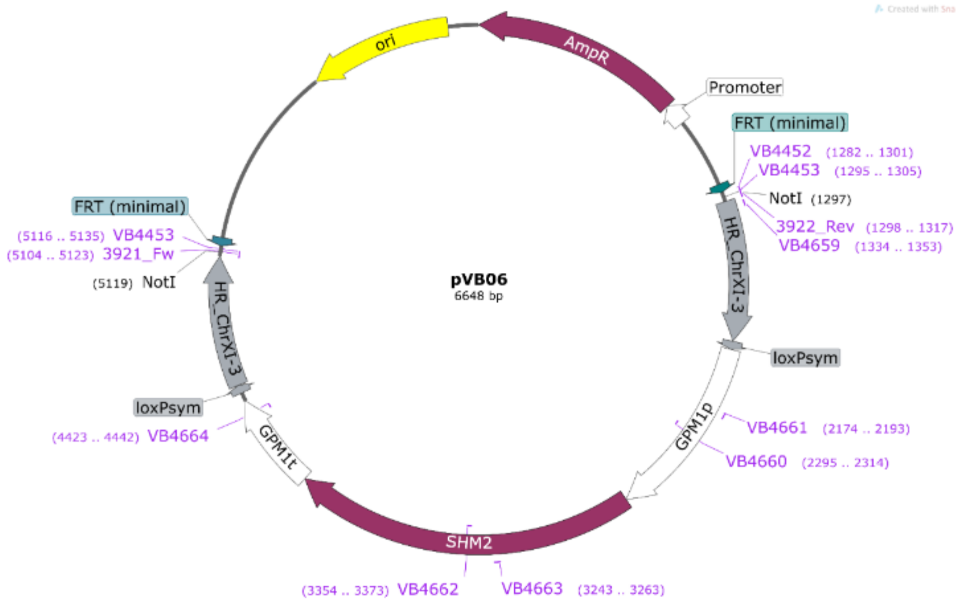


**Plasmid map 2:** Map of pVB06 plasmid with *SHM2* expression module under the *GPM1* promoter and terminator cloned between ChrXI-3 integrable homology arms.

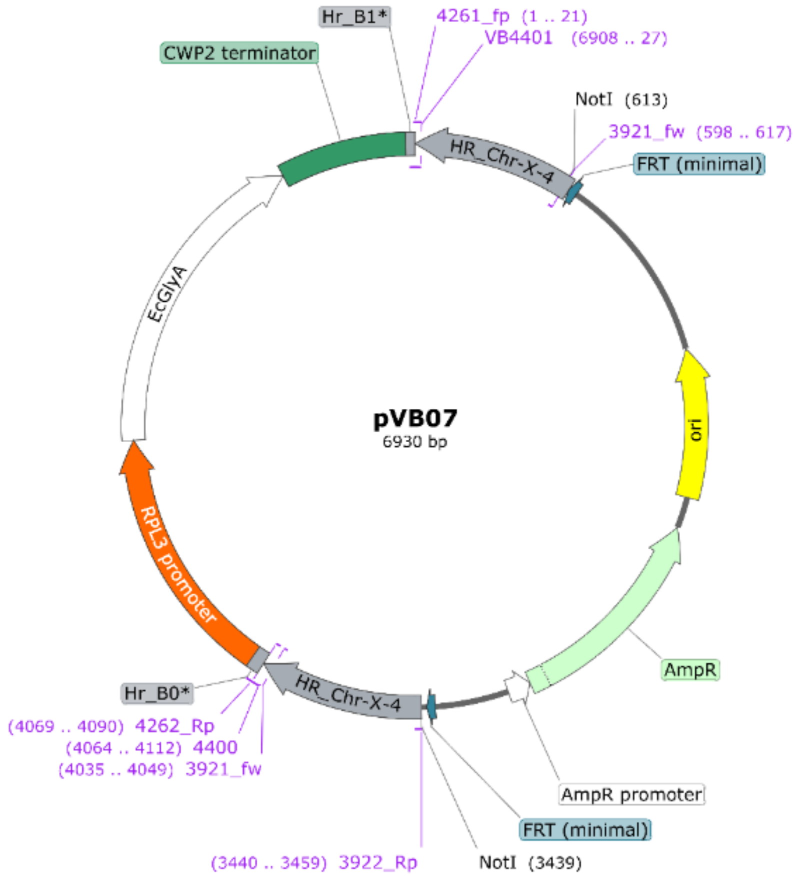


**Plasmid map 3:** Map of pVB07 plasmid with *EcGlyA* expression module under the *RPL3* promoter and *CWP2* terminator cloned between ChrX-4 integrable homology arms.

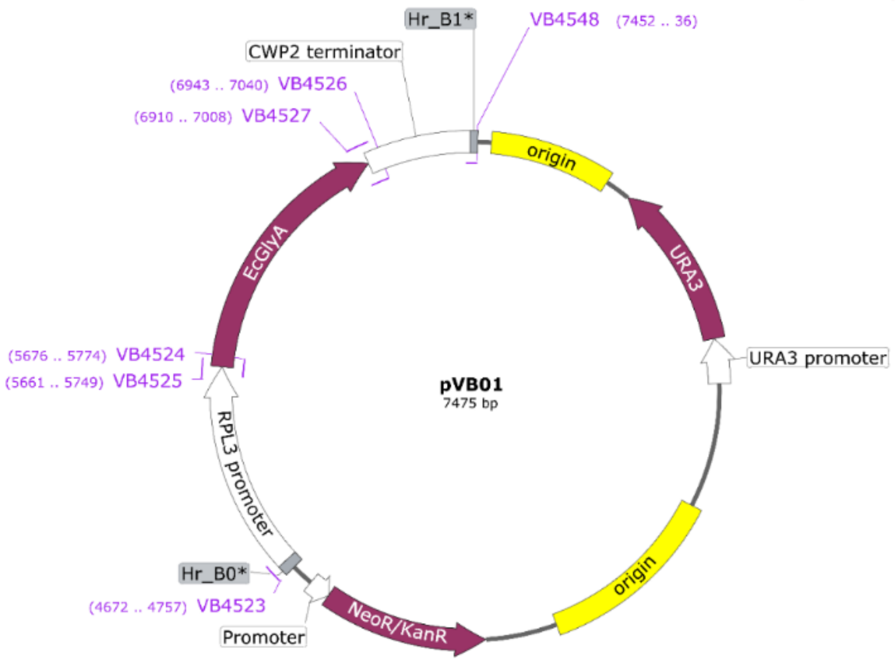


**Plasmid map 4:** Map of pVB01 plasmid with *EcGlyA* expression module under the *RPL3* promoter and *CWP2* terminator.

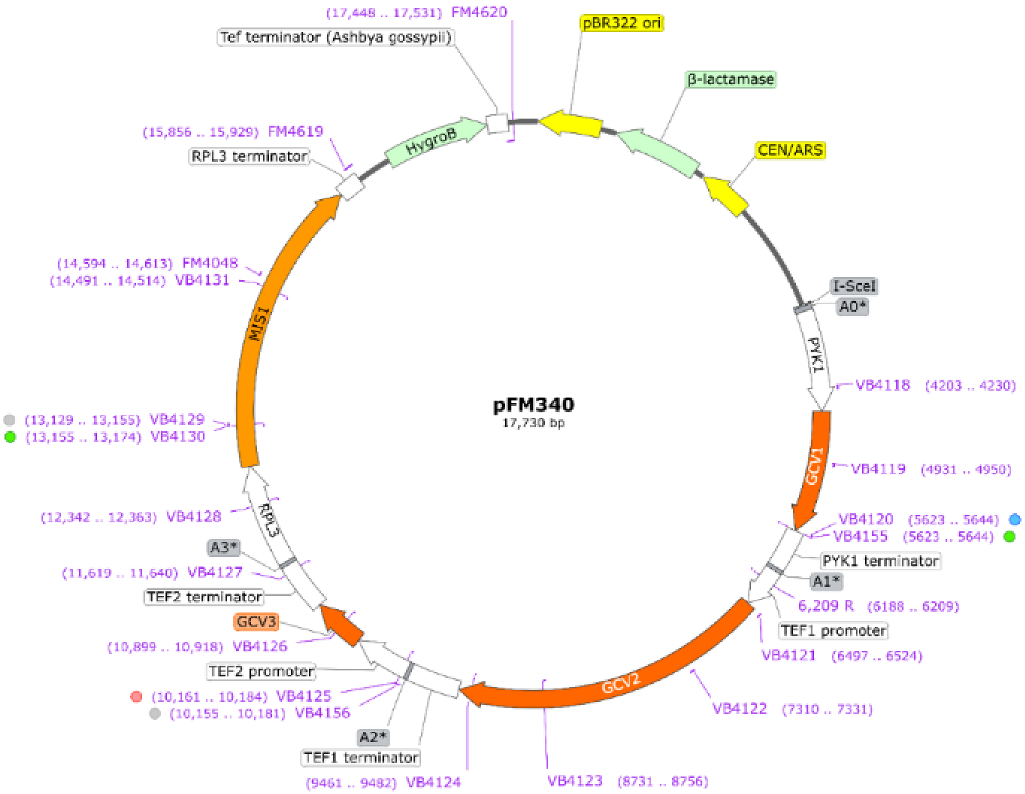


**Plasmid map 5:** Plasmid map of pFM340 with *MIS1*, *GCV1*, *GCV2*, and *GCV3* genes under the promoter and terminators of *PYK1*, *TEF1*, *TEF2*, and *RPL3,* respectively. pFM340 carries a Hygromycin resistance cassette for selection in the yeast system. It is a centromere plasmid with CEN/ARS Ori.

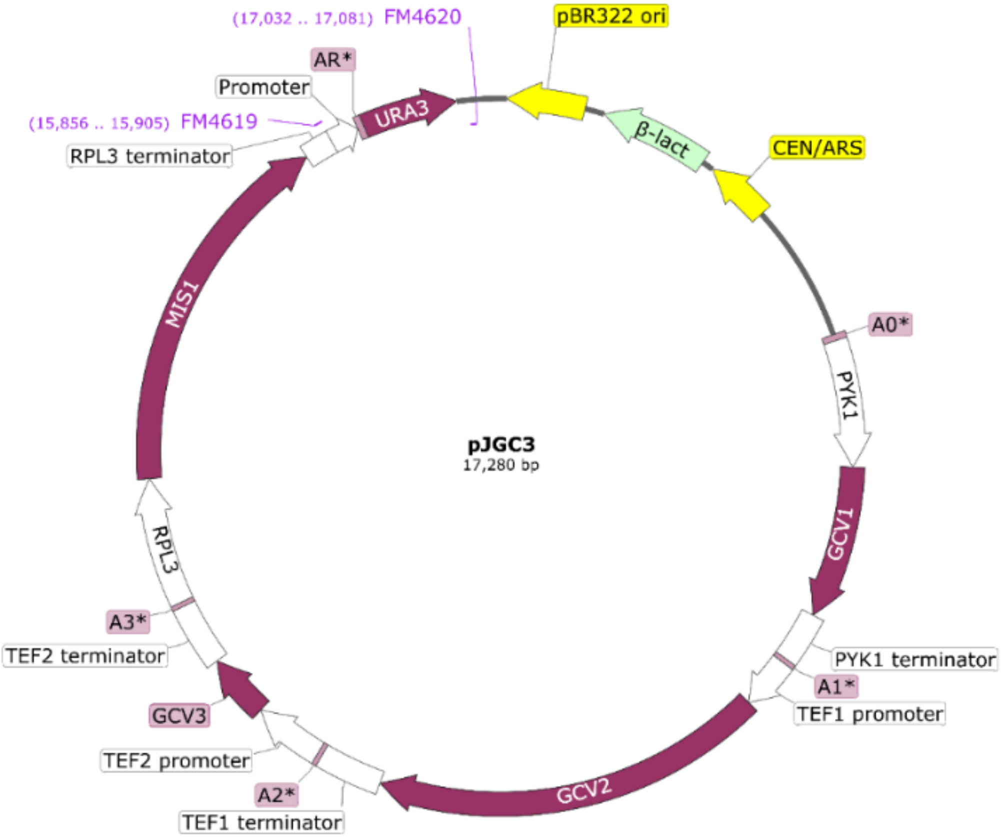


**Plasmid map 6:** Plasmid map of pJGC3 with *MIS1*, *GCV1*, *GCV2*, and *GCV3* genes under the promoter and terminators of *PYK1*, *TEF1*, *TEF2*, and *RPL3,* respectively. pFM340 carries a *URA3* selection marker to maintain in the yeast system. It is a centromere plasmid with CEN/ARS Ori.

### Supplementary tables

**Supplementary table 1: List of yeast strains and expression modules of the RGP used in this study**

| Name | Strains | Genotype | Glycine synthesis module overexpression | Carrier | SHMT overexpression module | Carrier |
|---|---|---|---|---|---|---|
| **VBS01** | S288c_ΔS | S288c *agx1, gly1, ser1* | No | — | No | — |
| **VBS03** | S288c_ΔS-pCfB-Cas9-Kan$^R$ | S288c *agx1, gly1, ser1* | No | pCfb2312 | No | pCfb2312 |
| **VBS04** | S288c_ΔS-ChrX-4::SHM1 | S288c *agx1, gly1, ser1* | No | — | *SHM1* | ChrX-4 |
| **VBS05** | S288c_ΔS-ChrXI-3::SHM2 | S288c *agx1, gly1, ser1* | No | — | *SHM2* | ChrXI-3 |
| **VBS06** | S288c_ΔS-ChrX-4::EcGlyA | S288c *agx1, gly1, ser1* | No | — | *EcSHMT (GlyA)* | ChrX-4 |
| **VBS08** | S288c_ΔS-ChrX-4::SHM1-pFM340-Hyg$^R$ | S288c *agx1, gly1, ser1* | *GCV1, GCV2, GCV3, MIS1* | pFM340 | *SHM1* | ChrX-4 |
| **VBS09** | S288c_ΔS-ChrXI-3::SHM2-pFM340-Hyg$^R$ | S288c *agx1, gly1, ser1* | *GCV1, GCV2, GCV3, MIS1* | pFM340 | *SHM2* | ChrXI-3 |
| **VBS10** | S288c_ΔS-ChrX-4::EcGlyA-pFM340-Hyg$^R$ | S288c *agx1, gly1, ser1* | *GCV1, GCV2, GCV3, MIS1* | pFM340 | *EcSHMT* (GlyA) | ChrX-4 |
| **VBS18** | VBS10_Δgdh1::108bp | S288c *agx1, gly1, ser1, gdh1* | *GCV1, GCV2, GCV3, MIS1* | pFM340 | *EcSHMT* (GlyA) | ChrX-4 |
| **VBS19** | S288c_WT_Δgdh1::108bp | S288c *gdh1* | No | — | No | — |

**ΔS**-serine biosensor strain; **Chr**-chromosome; **SHMT**-serine hydroxymethyltransferase; **Ec**-*E. coli*.

**Supplementary table 2: List of plasmids used in this study:** Chr-**chromosome;**

| Plasmid | Details | Used for | Source |
|---|---|---|---|
| pWS082 | sg entry vector | Gene deletion | Addgene # 90516 |
| pWS173 | Cas9-Kan$^R$ | Gene deletion | Addgene # 90960 |
| pWS174 | Cas9-Nat$^R$ | Gene deletion | Addgene # 90961 |
| pWS175 | Cas9-Hyg$^R$ | Gene deletion | Addgene # 90962 |
| pCfB2312 | Cas9-Kan$^R$ | Pathway integration | Addgene Kit # 1000000098 |
| pCfB3041 | Nat$^R$-sgChrX-3 | Target-ChrX-3 | Addgene Kit # 1000000098 |
| pCfB3042 | Nat$^R$-sgChrX-4 | Target-ChrX-4 | Addgene Kit # 1000000098 |
| pCfB3045 | Nat$^R$-sgChrXI-3 | Target-ChrXI-3 | Addgene Kit # 1000000098 |
| pCfB3034 | HrChrX-3 | ChrX-3 donor | Addgene Kit # 1000000098 |
| pCfB3035 | HrChrX-4 | ChrX-4 donor | Addgene Kit # 1000000098 |
| pCfB2904 | HrChrXI-3 | ChrXI-3 donor | Addgene Kit # 1000000098 |
| pVB05 | PGK1p-SHM1ox | SHM1-ChrX-4 donor | In this study |
| pVB06 | GMP1p-SHM2ox | SHM2-ChrXI-3 donor | In this study |
| pVB07 | RPL3p-EcGlyAox | EcGlyA-ChrX-4 donor | In this study |
| pFM340 | Hyg$^R$-MIS1-GCV1-3ox | Pathway expression | In this study |

**Supplementary table 3: List of strains and media conditions used for testing glycine and serine synthesis using VBS08, VBS09, and VBS10 in SC-Gly-Ser and SM media conditions: SC:** Synthetic Complete or semi-rich medium, **SM:** Synthetic minimal medium. **DO-Gly-Ser** is a mix of all the amino acids nucleotides in recommended quantities for optimal growth of yeast except serine and glycine, (**DO:** dropout);

| Media → | SM =1x YNB, 100 mM $(NH_4)_2SO_4$, 100 mM glucose | | | SC=1x YNB, DO-Gly-Ser, 100 mM $(NH_4)_2SO_4$,100 mM glucose | | |
|---|---|---|---|---|---|---|
| Strains ↓ | Growth positive control | Test glycine and serine synthesis | Growth negative control | Growth positive condition | Test glycine and serine synthesis | Growth negative control |
| VBS08 | SM, Glycine, formate | SM, formate | SM | SC-Ser+Gly formate | SC-Ser-Gly, formate | SC-Ser-Gly |
| Growth | ++ | - | - | ++ | - | - |
| VBS09 | SM, Glycine, formate | SM, formate | SM | SC-Ser+Gly formate | SC-Ser-Gly, formate | SC-Ser-Gly |
| Growth | ++ | - | - | ++ | - | - |
| VBS10 | SM, Glycine, formate | SM, formate | SM | SC-Ser+Gly formate | SC-Ser-Gly, formate | SC-Ser-Gly |
| Growth | ++ | - | - | ++ | + | - |

**Supplementary table 4: List mutations identified from the NGS data VBS10 strains during ALE**

| Identified mutations in the coding regions | | | | | | | |
|---|---|---|---|---|---|---|---|
| Gene | Mutation | Ev02 | Ev06 | Ev16 | Ev21 | Annotation | Description |
| GDH1 ← | Δ108 bp | + | + | + | + | coding (223-330/1365 nt) | glutamate dehydrogenase (NADP(+)) GDH1 |
| UTP10 ← | C→A | - | + | + | + | M1I (ATG→ATT) † | snoRNA-binding rRNA-processing protein UTP10 |
| PET9 ← | A→G | - | + | + | + | I275T (ATT→ACT) | ADP/ATP carrier protein PET9 |
| IDH1 ← | G→T | - | + | + | + | P299Q (CCA→CAA) | isocitrate dehydrogenase (NAD(+)) IDH1 |
| SUI2 | G→T | - | - | - | + | R88L (CGT→CTT) | translation initiation factor eIF2 subunit alpha |
| ASH1 | Δ3 bp | - | - | - | + | coding (1237-1239/1767 nt) | DNA-binding transcription repressor ASH1 |

| Identified mutations in the intergenic regions | | | | | | | |
|---|---|---|---|---|---|---|---|
| Mutation | Ev02 | Ev06 | Ev16 | Ev21 | Annotation | Gene | Description |
| A→G | + | + | + | + | intergenic (-100/-807) | *GSH1 ← / → LSB6* | glutamate--cysteine ligase/1- phosphatidylinositol 4-kinase LSB6 |
| C→A | + | + | + | + | intergenic (-504/-184) | *MET6 ← / → IES5* | 5-methyltetrahydropteroyltriglutamate- homocysteine S-methyltransferase/Ies5p |
| $(C)_{5\rightarrow4}$ | - | + | + | + | intergenic (+4733/-3179) | *VAR1 → / → 21S_RRNA* | mitochondrial 37S ribosomal protein VAR1/21S ribosomal RNA (Ori) |
| $(A)_{13\rightarrow14}$ | + | + | + | - | intergenic (-240/+391) | *ERG26 ← / ← EFM5* | sterol-4-alpha-carboxylate 3-dehydrogenase (decarboxylating)/protein-lysine N-methyltransferase |
| $(A)_{11\rightarrow12}$ | + | - | - | - | intergenic (+264/-4749) | *COX3 → / → tM(CAU)Q2* | cytochrome c oxidase subunit 3/tRNA-Met |

**Supplementary table 5: List of components used in DO-Ser-Gly mix.**

| **S1 Table: Composition of Drop-Out mix used in SC medium** | | | |
|---|---|---|---|
| **S. No.** | **Chemical component** | **mg/L** | **SC-Ser-Gly** |
| 1 | Adenine | 18 | + |
| 2 | p-Aminobenzoic acid | 8 | + |
| 3 | Leucine | 380 | + |
| 4 | Alanine | 76 | + |
| 5 | Arginine | 76 | + |
| 6 | Asparagine | 76 | + |
| 7 | Aspartic acid | 76 | + |
| 8 | Cysteine | 76 | + |
| 9 | Glutamic acid | 76 | + |
| 10 | Glutamine | 76 | + |
| 11 | Glycine | 76 | - |
| 12 | Histidine | 76 | + |
| 13 | Myo-inositol | 76 | + |
| 14 | Isoleucine | 76 | + |
| 15 | Leucine | 76 | + |
| 16 | Lysine | 76 | + |
| 17 | Methionine | 76 | + |
| 18 | Phenylalanine | 76 | + |
| 19 | Proline | 76 | + |
| 20 | Serine | 76 | - |
| 21 | Threonine | 76 | + |
| 22 | Tryptophan | 76 | + |
| 23 | Tyrosine | 76 | + |
| 24 | Uracil | 76 | + |
| 25 | Valine | 76 | + |

**Note:** Glycine and serine are excluded from the mix.

**Supplementary table 6: List of the primers used for screening and sequencing**
Other primers used for PCRs and cloning were directly indicated on the respective plasmid maps 1-6.

| Table S11 | List of primers used in this study | | | |
|---|---|---|---|---|
| S. No | Name | Sequence | Purpose | Target |
| 1 | **ASK-FN2** | CGGCCTTTTTACGGTTCCTG | Seq. sgOligo insertion | Sequencing |
| 2 | **3569** | **gacttt**GTTGTCGCAAAATAGCTATG | sgRNA | AGX1_pWS_f |
| 3 | **3570** | **aaac**CATAGCTATTTTGCGACAAC**aa** | | AGX1_pWS_r |
| 4 | **3571** | gactttGGTCTCAAACTACATTGTGA | | Gly1_pWS_f |
| 5 | **3572** | aaacTCACAATGTAGTTTGAGACCaa | | Gly1_pWS_r |
| 6 | **3573** | gactttCCGGTGACTAAATAACCGGC | | SER1_pWS_f |
| 7 | **3574** | aaacGCCGGTTATTTAGTCACCGGaa | | SER1_pWS_r |
| 8 | **3634** | TGTCTCCGGTTAGTGTGTGC | Knockout screening | AGX1_f |
| 9 | **3635** | GCCCCAGAAGTGCTTTTTGG | | AGX1_r |
| 10 | **3636** | TACACACGGCCCCAAATTGT | | GLY1_f |
| 11 | **3637** | CACCGATCGAGGAAGCCTTT | | GLY1_r |
| 12 | **3638** | AAAAAGCGTTTCGTGGAGCA | | SER1_f |
| 13 | **3639** | GCTCCCTCTTGCACTTCACT | | SER1_r |
| 14 | **3632** | ATCAACAACAGAGGACATATGCC | sgRNA expression cassette amplificaiton | pWS082_f |
| 15 | **3633** | ATCCTGCACTCATCTACTACCC | | pWS082_r |
| 16 | **4415** | gaagtgccattccgcctgacct | TWIST adapter primers for donor DNA amplificaiton | Donor DNA amplificaiton |
| 17 | **4416** | cactgagcctccacctagcct | | |
| 18 | **4619** | CTACATACATGTACATATATTTAAACATGTAAACCCGTCCATTATATTGCGACATGGAGGCCCAGAATACCCTC | pFM0340_assembly | Plasmid assembly |
| 19 | **4620** | TGAATAACTGCTCCCGTTGTAAAAGTATCCGTAGTGTGACTGAAACCTTACAGTATAGCGACCAGCATTCACATACGATTGACG | pFM0340_assembly | |
| 20 | **4656** | AGTGACCACCATCTGGCAAA | pCfB2904 SHM1 seq R2 | Sequencing donor pathway modules |
| 21 | **4657** | ACATCTGCATAATAGGCATTTGCA | pCfB2904 SHM1 seq F3 | |
| 22 | **4658** | AAACACTTCCTTTTTCTGGCCC | pCfB2904 SHM1 seq R3 | |
| 23 | **4659** | CGTTTGCGCATCCTCTTTTT | pCfB3042 SHM2 seq F1 | |
| 24 | **4660** | AGACGAAGCTTGTGTGTGGG | pCfB3042 seq R1 | |
| 25 | **4661** | GCGCGATGGTGCTAAGTTAC | pCfB3042 seq F2 | |
| 26 | **4662** | GCGGCGATCAAACCAGAAAT | pCfB3042 seq R2 | |
| 27 | **4663** | CCAAAGGTTCTTGTTGCTGGT | pCfB3042 seq F3 | |
| 28 | **4664** | CGAATTCGCGGGGTGTACAT | pCfB3042 seq R3 | |
| 29 | **6643** | cccaaagctaagagtcccattttattc | pCFB3035 donor X-4 Fw | pCfB donor amplificaiton |
| 30 | **6644** | ctggtgaggatttacggtatgatcatg | pCFB3035 donor X-4 Rev | |
| 31 | **3921** | GGCCGCGCTGAGGTCTTAAT | Easy_clone_donor_fw | |
| 32 | **3922** | GGCCGCGCTGAGGGTTTAAT | Easy_clone_donor_rev | |
| 33 | **4401** | gccagggttttcccagtcacgacgttgtgtttcacaaccagagattcagg | EcGlyA expression cassette | Fw primer |
| 34 | **4402** | gccagggttttcccagtcacgacgttgTACGCAGTCTTCGGGTAGTAAAATAG | EcGlyA expression cassette | Rev primer |
| 35 | **4048** | GGACGCATGGTTATTGGTGC | pFM_0274_f | Plasmid assembly |
| 36 | **4049** | GATCGACCTTCCATGGGGTC | pFM_0274_r | |
| 37 | **4400** | gcgtgcgatgagcgacctcatgctatataaggtttcagtcacactacgg | pVB07 assembly | |
| 38 | **4401** | gccagggttttcccagtcacgacgttgtgtttcacaaccagagattcagg | pVB07 assembly | |
| 39 | **6938** | gactttGCCAAGGGTCCATACAAGGG | sgRNA | sgGDH1 |
| 40 | **6939** | aaacCCCTTGTATGGACCCTTGGCaa | | |
| 41 | **6940** | gactttACACATAGACCACCCTTGTA | | |
| 42 | **6941** | aaacTACAAGGGTGGTCTATGTGTaa | | |
